# Investigating the skin response of Striped Catfish to acute ammonia stress reveals a potential exosome-based ammonia excretion system

**DOI:** 10.64898/2026.01.07.696813

**Authors:** Wei-Hsuan Hsiao, Shun-Yi Lin, Ying-Hsien Wu, Han-Chen Ho, Tsung-Lin Liu, Liang-Chun Wang

## Abstract

Ammonia can rapidly accumulate in intensive aquaculture and pose serious threats to fish health. Fish skin functions as a vital immune-related organ and a significant surface barrier against ammonia; however, little is known about the skin’s response and to acute ammonia stress. In this study, Striped catfish were exposed to a half-lethal concentration of ammonia to evaluate skin responses. We found generalized inflammation in the liver, spleen, and skin. However, the skin tissue showed subtle signs of damage, with tissue ammonia concentration accumulating and then decreasing. Further transcriptomic analysis of skin gene function suggested a hypothesized ammonia regulation occurring within cell compartments. Cellular examination by transmission electron microscopy further showed that accumulated exosome-suspecting vesicles near the epithelial surface were associated with ammonia regulation during acute ammonia stress. Using a newly established Striped catfish skin epithelial cell line, along with an exosome inhibitor, confirms the presence of exosome vesicles and their role in ammonia excretion. In conclusion, acute ammonia stress induces inflammation in Striped catfish skin, while active ammonia excretion via exosomes may alleviate ammonia accumulation, thereby maintaining skin tissue integrity. This study explores Striped catfish skin responses to acute ammonia stress and highlights a newly identified mechanism of ammonia regulation in the skin.

**Highlights:**

- Fish skin can regulate ammonia concentrations and stress.
- Exosomal vesicles are involved in the excretion of ammonia from the skin.

## 1. Introduction

Striped catfish, with rapid growth, is one of the most economically important fish species in eastern Asia [1]. Striped catfish farming is typically intensive, which also entails substantial risks, including chemical and biological hazards [2, 3]. Among chemical hazards, nitrogen-based pollution, often referred to as ammonia nitrogen, is most prevalent. Ammonia nitrogen, often referred to as total ammonia nitrogen (TAN) [4], exists in the combination of ionic (NH_4_^+^) and nonionic (NH_3_) forms. TAN mainly comes from feed residues, fish excretion, and dead fish [5–7]. The concentration of TAN can vary widely and rapidly [8]. For example, a study reported that TAN concentrations ranged from 1.4 to 10 mg/L, with the highest concentration recorded in a collapsible pond and the lowest in a natural pond [9]. However, TAN concentrations in intensive aquaculture farming systems, such as shrimp ponds, can reach 46 mg/L in 14 days [10]. TAN are toxic to aquatic animals. In fish, ammonia can diffuse through the gills and accumulates in tissues [11], leading to reduced growth, activity reduction, cell death, metabolic disorder, oxidative stress, and immune responses [12–17]. In addition, oxidative stress, inflammation, and apoptosis increased significantly in Yellow catfish *P. fulvidraco* under 57 mg/L TAN stress for 96 hr [18]. Therefore, ammonia-induced damage to catfish and the production needs associated with it is required to be addressed.

Collective studies have shown that fish can manage the impact of ammonia through ammonia detoxification and immune modulation. Ammonia detoxification is often referred to as reducing the intracellular ammonia concentration. One way to transport ammonia out of a cell is through active transportation through Rhesus (Rh) glycoproteins [19]. Mud loach *P. dabryanus* significantly increases the Rh glycoproteins expression under acute 420 mg/L TAN stress for 48 hr [20]. Another way is to metabolize the ammonia through amino acid metabolism [21]. Magur catfish *C. magur* increases the aminoa acid metabolism under 350 mg/L TAN stress for 24 hr [22]. While decreasing ammonia concentration, the immune system, more specifically, inflammation, also helps cope with acute ammonia stress and cell damage, thereby maintaining cell survival [23, 24]. The instant inflammation can urgently prevent damage and unnecessary energy consumption [23]. Examples of Yellow catfish *P. fulvidraco* under acute ammonia stress increase the inflammatory cytokine expression in the liver under acute 125 mg/L TAN stress for 48 hr [25]. While acute ammonia stress can be managed by acute ammonia excretion, metabolic transformation, and immune responses, chronic ammonia stress can still lead to increased tissue accumulation and eventually immunosuppression in major internal organs [26]. For example, the innate immunity of spleen and head kidney in Wuchang bream *M. amblycephala* were significantly inhibited under chronic 30 mg/L TAN stress for 30 days [16]. However, different from internal organs [27], fish skin was reported to be unaffected or minimally changed under ammonia exposure [28]. The findings raise the question of how the skin copes with ammonia and whether it may employ alternative protective mechanisms.

Fish skin, together with skin mucus, is a metabolically active and multifunctional organ involved in immunity, excretion, and osmoregulation [29]. The skin-associated lymphoid tissue (SALT) is part of the fish immune system in peripheral immune organs [30]. Skin contains diverse leukocytes, plasma cells, macrophages, and granulocytes, and has been shown to excrete nitrogenous waste [31, 32]. However, under high ammonia concentration, ammonia can be seen accumulated in the skin [33, 34]. Even though, when comparing with liver, gill, and spleen, fish skin shows minimal histological alterations under ammonia stress, suggesting a higher recovery capacity [35]. In European seabass *D. labrax* under 11.90 mg/L TAN stress for 48 hr, skin thickness remained the same with no cell loss but only subtle morphology alteration [28]. With these findings, the potential ammonia excretion, its alternatives, and the intertwined immunometabolic responses of fish skin to ammonia are of interest. Investigating the responses of fish skin to acute ammonia stress is the first step and thus important for mechanistically explaining ammonia management.

Given the limited research on fish skin responses to acute ammonia stress, the present study aims to investigate how fish skin responds to and regulates ammonia stress through an acute ammonia challenge. The obtained responses will provide insights not only into long-term ammonia management in fish but also into fish-farming strategies during elevated ammonia exposure and into aquaculture management practices.

## 2. Materials and methods

### 2.1. Fish husbandry

Striped catfish (*Pangasianodon hypophthalmus*) purchased from a local aquaculture vendor were reared in 150 L husbandry tanks and maintained a constant photoperiod (12 hr light; 12 hr dark) along with sufficient aeration in the aquaculture room at the Department of Marine Biotechnology and Resources. Fish grown to an average length of 10.0±1 cm were used for the experiment. Tap water was used in this study and routinely monitored during husbandry. Temperature (26.26±0.26 °C) and pH (8.26±0.17) were measured using a pH meter (Suntex Instruments, Taiwan). Dissolved oxygen concentrations (6.36±0.40 mg/L) were measured using a YSI 550A meter (YSI, USA). TAN (<0.5 mg/L) was measured using a Hach DR1900 Spectrophotometer (Hach, USA). The Institutional Animal Care and Use Committee (IACUC) of National Sun Yat-sen University permitted the experiment with approval number 11104.

### 2.2. Experimental design and sample collection

NH_4_NH_3_ (analytical purity, 99.5% purity) was used to represent soluble ammonia nitrogen, and a pretest of the toxicity of Striped catfish was performed before the formal test. The TAN concentration of 40 mg/L was determined in this study based on our preliminary experiment on the half-lethal concentration (Fig. S1). The non-ammonia control group and ammonia group were established in 40 L of fish tank water, each with six fish. The photoperiod was maintained at 12 hr of light and 12 hr of darkness. Furthermore, the hypobromite oxidation method was used to measure ammonia nitrogen concentration in the aquaculture water every 6 hr to confirm a constant ammonia concentration. The pH was kept constant at 8.26±0.17. Feeding was stopped during the experiment. Water quality was managed in the same manner as during the rearing period. Samples were collected at 0, 3, 6, 12, and 24 hr. At each time point, the experimental fish were randomly selected from the fish tank and anesthetized with MS-222 (150 mg/L) buffered with sodium bicarbonate. Skin tissues were gently removed, frozen in liquid nitrogen, and stored at -80 °C until further analysis.

### 2.3. Physiological evaluation

Fish health deterioration was evaluated using a video-based scoring system adapted from Jarvis et al. and Kartikaningsih et al. [36, 37], based on seven behavioral traits (Table 2), each scored from 0 to 5 by at least three observers. Scores were summed and converted into percentages to represent health deterioration. Photos of fish in each group were taken before sampling, and ImageJ was utilized to evaluate the visual fin and skin damage. Fin damage was quantified by comparing the area and intensity of red coloration to that of normal fin tissue. Skin damage was quantified by comparing the areas of redness, hemorrhage, discoloration, and ulceration to those of adjacent normal skin areas [38, 39].

### 2.4. Histological examination of the skin

Tissue samples were fixed in Bouin’s solution for 48 hr. Subsequently, the fixed samples were dehydrated with alcohol and Histo-Clear II (National Diagnostics, USA), then embedded in paraffin wax. Samples were sectioned into 7-10 µm sections using a Histocut Rotary Microtome (Leica Model 820, REICHERT, USA). The paraffin was removed from the tissue using Histo-Clear II and a graded series of alcohol. The tissue was then rehydrated and stained with hematoxylin and eosin staining with or without Nessler’s reagent.

### 2.5. RNA extraction and Illumina sequencing

Total RNA of skin tissues was extracted using TRIzol Reagent (Thermo Fisher Scientific, USA) according to the manufacturer’s instructions. Purified RNA was dissolved in nuclease-free water (Promega, USA). The DeNovix DS-11 Spectrophotometer (DeNovix, USA) was used to determine the concentration and quality of purified RNA. RNA sequencing was performed using the Illumina sequencing platform by Genomics BioSci. & Tech ( Genomics BioSci. & Tech, Taiwan). mRNA was purified using poly-T oligo-attached magnetic beads and subsequently fragmented for cDNA synthesis. First-strand cDNA was synthesized using reverse transcriptase and random primers, followed by second-strand synthesis incorporating dUTP to retain strand specificity. A single ‘A’ nucleotide was then added to the 3′ ends of the double-stranded cDNA, and indexing adapters were ligated to both the 5′ and 3′ ends. Adapter-ligated fragments were selectively amplified by PCR to enrich for properly constructed libraries. The quality and concentration of the resulting libraries were validated using an Agilent 2100 Bioanalyzer (Agilent Technologies, USA) and a real-time PCR system. Finally, the six libraries were sequenced on the Illumina NovaSeq 6000 platform (Illumina, USA) to generate paired-end reads.

### 2.6. Quality control and mapping

The raw data filtration included removing reads with adapters, reads containing too many Ns, and low-quality reads (Qphred ≤20). At the same time, Q20, Q30, and GC content were calculated for the clean data. All subsequent analyses were high-quality analyses based on clean data. *Pangasianodon hypophthalmus* reference genome data were obtained from NCBI (fPanHyp1.pri). The clean reads were aligned to the reference genome using STAR [50] to obtain their positions relative to the reference genome.

### 2.7. Analysis of differentially expressed genes (DEGs) and functional enrichment

The abundance of each transcript region was quantified as Counts Per Million (CPM) values. These CPM values were subsequently normalized using the Trimmed Mean of M-values (TMM) method to adjust for variations in library size and composition across samples. This normalization step is crucial for accurate downstream differential expression analysis. Differential expression analysis of RNAs, including both mRNAs and lncRNAs, was conducted to identify transcripts with statistically significant changes in expression between experimental conditions. The Limma R package [51] was used to compare two distinct ammonia groups. For pairwise comparisons across multiple groups or conditions, the edgeR R package [40] was utilized. Transcripts were identified as differentially expressed if they exhibited an absolute log2 Fold Change (|log2FC|) of ≥ 0 and an adjusted P-value (Benjamini-Hochberg) of ≤ 0.05. This P-value threshold ensures a controlled false discovery rate (when using adjusted P-values) or a controlled Type I error rate. To elucidate the potential biological functions and pathways associated with the identified DEGs, Gene Ontology (GO) enrichment analysis and Kyoto Encyclopedia of Genes and Genomes (KEGG) pathway analysis were performed using the R package clusterProfiler [41]. GO analysis aimed to identify over-represented biological processes, molecular functions, and cellular components, using the resources available at the Gene Ontology Consortium [42–44]. KEGG pathway analysis was conducted to map DEGs to relevant metabolic and signaling pathways, using the KEGG database [45, 46] accessed via clusterProfiler. For both GO and KEGG enrichment analyses, gene sets, or pathways were considered significantly enriched if the associated adjusted P-value (Benjamini-Hochberg) was < 0.05. This threshold allowed for the identification of key biological themes and regulatory networks perturbed under the investigated conditions.

### 2.8. Quantitative real-time PCR

cDNA was synthesized using 2 μg of total RNA with M-MLV Reverse transcriptase (Promega, USA) according to the manufacturer’s instructions. Five primer pairs of specific immune markers for real-time PCR were selected based on the National Center for Biotechnology Information (NCBI) Primer BLAST (Table 1). Basically, real-time PCR was performed using the GoTaq qPCR Master Mix (Promega, USA) on a Mic qPCR Cycler (Bio Molecular Systems, Australia). The thermal cycling profile consisted of an initial denaturation at 95 °C (for 2 min), followed by 40 cycles of denaturation at 95 °C (3 s) and an appropriate extension temperature (60 °C, 30 s). An additional temperature ramping step was performed to produce melting curves of the reaction from 65 °C to 95 °C. The housekeeping gene EF1-α was used as the reference gene, and relative fold changes were calculated from cycle threshold (Ct) values obtained by real-time PCR. The 2^−ΔΔCT^ method was used to determine gene expression levels.

**Table 1.**
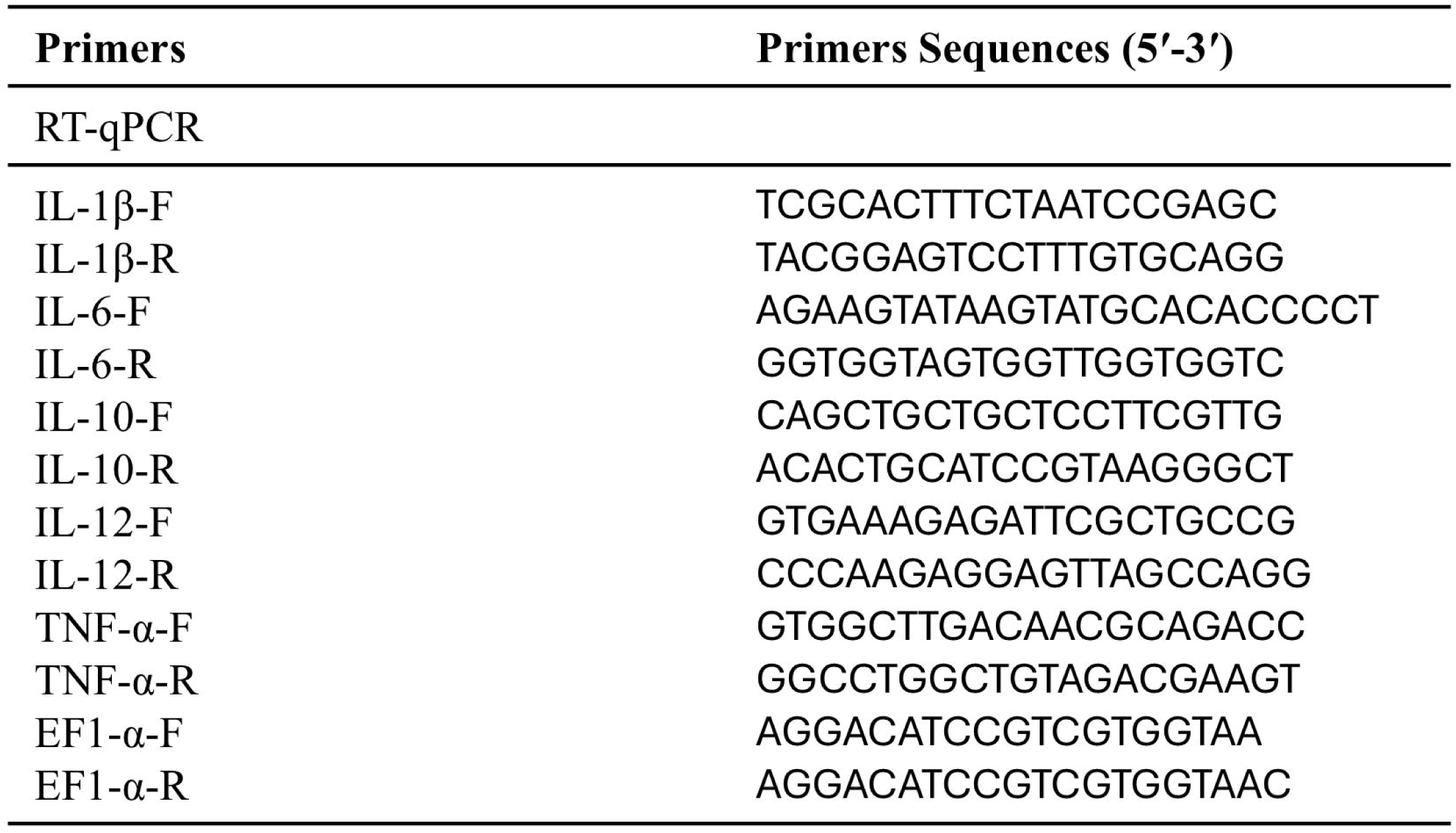
Immune cytokine gene primer.

### 2.9. Treatment of skin epithelial cells with an inhibitor, GW4869, and ammonia

Striped catfish skin epithelial cells [47] were plated in 12-well Cell Culture Plates (Alpha Plus, Taiwan) at 5 × 10^5^ cells/well, and treated with Leibovitz’s L-15 (Gibco™, USA) Medium with 20% exosome-free FBS for 24 hr. After 24 hr, GW4869 (Sigma-Aldrich, USA) was added to these cells at 10 μM for 1 hr, followed by a 6-hour challenge with 40 mg/L ammonia and recovery with or without GW4869 for 1 hr. Supernatants were collected after a 6 hr challenge and a 1 hr recovery for ammonia concentration analysis using the Nitrogen-Ammonia Reagent Set (Hach, USA). GW4869 was initially dissolved in DMSO (Fisher Scientific, USA) to a stock solution of 5 mM GW4869, then diluted into culture supernatant to achieve 10 μM. The final concentration of DMSO was smaller than 0.1%.

### 2.10. Statistical analysis

Statistical significance was assessed using the one-way analysis of variance (ANOVA) followed by Tukey’s honestly significant difference (HSD) multiple-comparison and Student’s unpaired t-test. All the data were confirmed to fit into a Gaussian distribution by a Shapiro-Wilk test for normality. All analyses were performed by GraphPad Prism 10 (GraphPad Software, La Jolla, CA).

## 3. Results

### 3.1. Striped catfish health deteriorates without visible skin damage under acute ammonia stress

To explore if Striped catfish, especially in skin, respond to acute ammonia stress in general, Striped catfish were divided into non-challenge controls and ammonia-challenged groups, and samples of each group were taken at 0, 6, 12, and 24 hr (Fig. 1A). Fish health deterioration was evaluated based on seven behavioral traits (Table 2) adapted from a scoring system established by Jarvis et al. and Kartikaningsih et al. [36, 37]. We found that the ammonia groups showed significant deterioration from 6 hr onward compared with non-challenge controls (Fig. 1B). Fish appearance is a physiological marker of tissue damage and health status under ammonia exposure [13]. To assess fish appearance, photographs of each group were taken to evaluate physiological damage [48]. We visually found a notable fin redness but no skin-based damage at 24 hr post-ammonia challenge (Fig. 1C). Quantitative analysis using ImageJ further confirmed that fin redness—expressed as the ratio of red area to normal fin area—was significantly increased from 6 hr and continued to intensify over time compared with non-challenged controls (Fig. 1D). In contrast, no visible lesions, damage, or significant alterations were observed in the skin at any given time point (Fig. 1E). These data suggest that acute ammonia stress casues health deteriorate but not skin damage in striped catfish.

**Figure 1.**
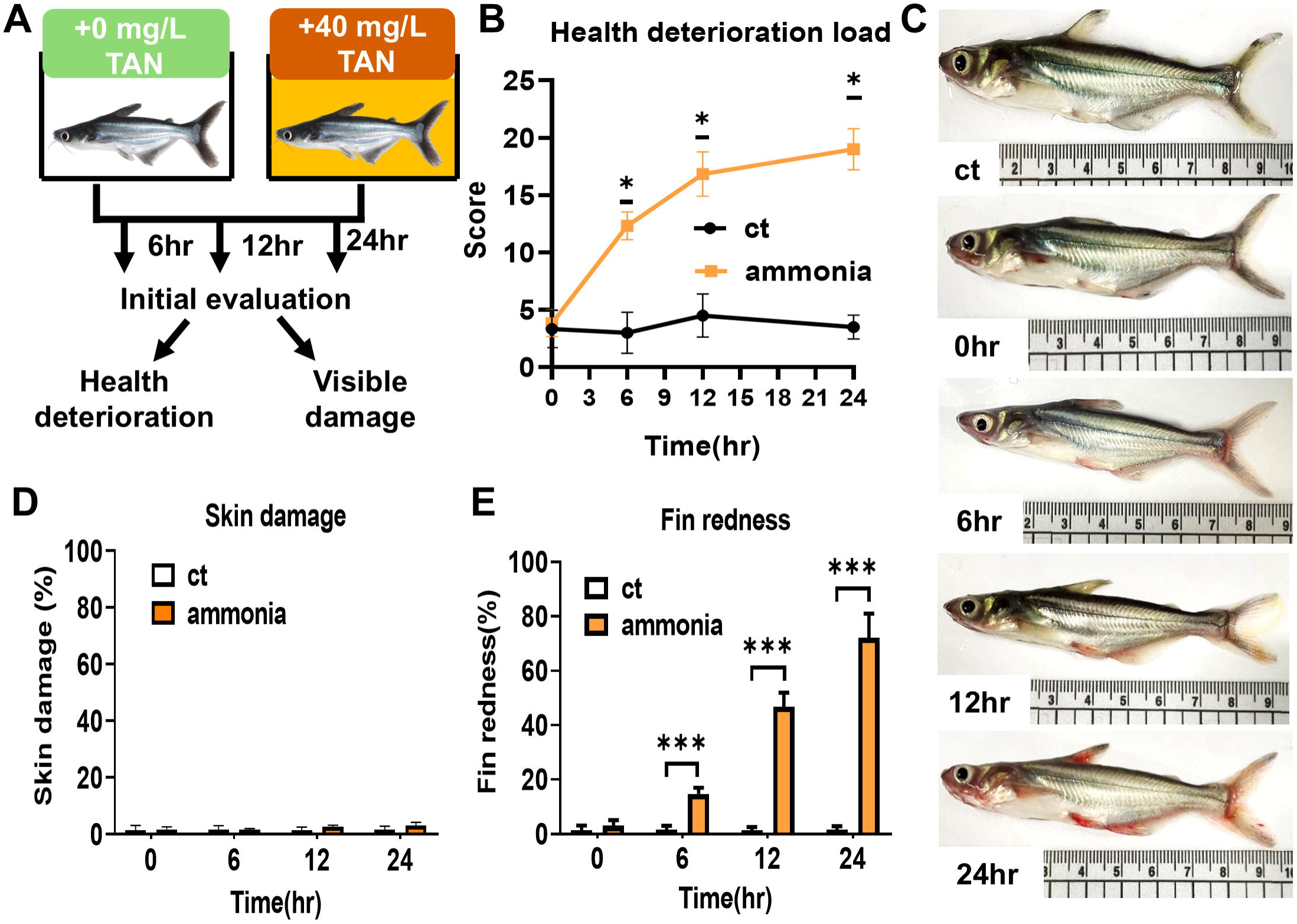
Assessment of visible deterioration of fish health over time under ammonia stress. Striped catfish challenged by 40 mg/L total ammonia nitrogen (TAN) were pictured before sampling. (A) Illustration of the experiment. (B) The health deterioration load was evaluated by video. Fish physiology (C) appearance, (D) fin redness, and (E) skin damage percentage were evaluated by ImageJ. Ct represents the non-challenge control group. Data are presented as mean ± SD. N = 6 fish in each group. All statistical significance was determined by Student’s unpaired t-test at * (p < 0.05), ** (p < 0.01), *** (p < 0.001).

**Table 2.** Parameters of health were evaluated in the health deterioration load of fish under ammonia challenge.

| Parameters of health evaluation |  |
| --- | --- |
| Standstill | Swimming sideways |
| Listless | Fin redness |
| Startled | Skin discoloration |
| Lesion |  |

### 3.2. Striped catfish skin expresses an increasing inflammatory response under acute ammonia stress

Inflammation is one of the hallmarks of ammonia-affected tissue. To further investigate how the skin was impacted by ammonia, expression of skin immune cytokines, including Interleukin 1 beta (IL-1β), Tumor Necrosis Factor Alpha (TNF-α), Interleukin 10 (IL-10), Interleukin 6 (IL-6), and Interleukin 12 (IL-12), was analyzed by qPCR. We found that IL-1β expression increased significantly at 6 hr and continued to rise thereafter (Fig. 2B). IL-12 and TNF-α expression were significantly increased at 12 hr and continued to rise thereafter (Fig. 2C & D). IL-6 and IL-10 expression, on the other hand, were significantly increased only at 24 hr (Fig. 2E & F). We also examined liver and spleen, the most commonly reported organs affected by ammonia, and found that IL-1β expression in these organs increased from 6 hr onward (Fig. S2A & B). Interestingly, the level of IL-1β in skin tissue was comparably higher than that in either the liver or the spleen. These data suggest that striped catfish skin tissue is affected by acute ammonia stress, even though no visual damage was observed.

**Figure 2.**
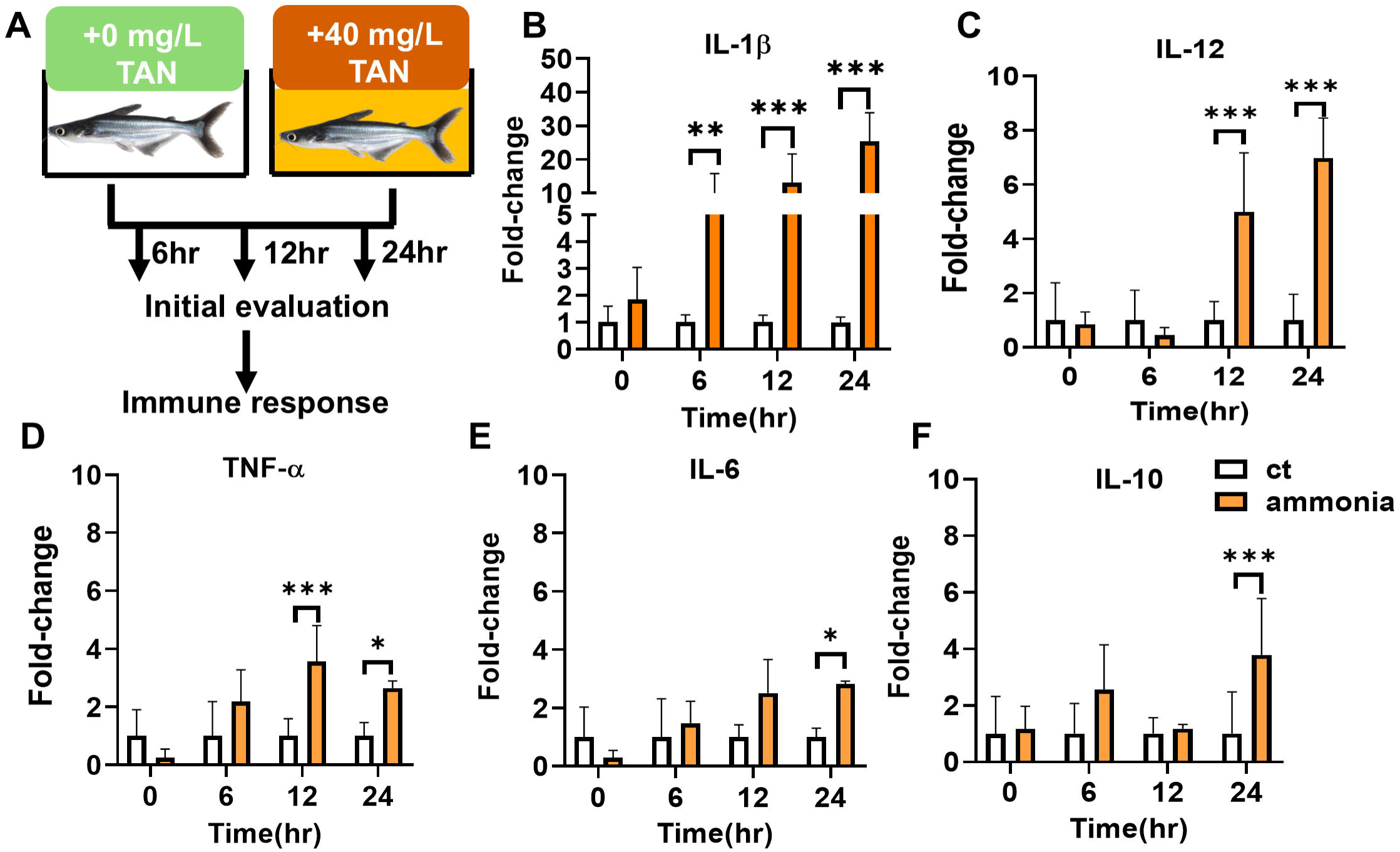
Evaluation of fish skin inflammation-related gene expression over time under ammonia stress. (A) Illustration of the experiment. (B to F) Expression levels of skin (B) IL-1β, (C) IL-12, (D) TNF-α, (E) IL-6, and (F) IL-10 were evaluated using qPCR. Ct represents the non-challenge control group. Data are presented as mean ± SD. N = 6 fish in each group. All statistical significance was determined by Student’s unpaired t-test at * (p < 0.05) ** (p < 0.01) *** (p < 0.001).

### 3.3. Striped catfish skin exhibits subtle damage at the tissue level

Inflammation triggered by noxious stimuli usually leads to tissue damage [49]. Since striped catfish skin shows inflammation without visible damage under acute ammonia stress, we hypothesized that damage may occur at the tissue and/or cellular levels. Hence, skin integrity at the tissue level was examined by paraffin-sectioned histology with hematoxylin and eosin (H&E) staining. We found no significant epidermal loss, yet observed a rough superficial surface with enlarged goblet cells and loss of membrane borders in club cells as the challenge progressed (Fig. 3A–D). To further assess whether reported epithelial loss in the epidermis, the thickness of the affected superficial and club cell layers was quantified. We found unchanged thickness in the superficial layer but a <20 μm reduction in club cell thickness (Fig. 3E & F), suggesting subtle epidermal thinning of the club cells from 12 hr onward (Fig. 3G). Taken together, these data indicate that, despite inflammation, skin tissue exhibits minimal epidermal damage at the tissue level under acute ammonia stress.

**Figure 3.**
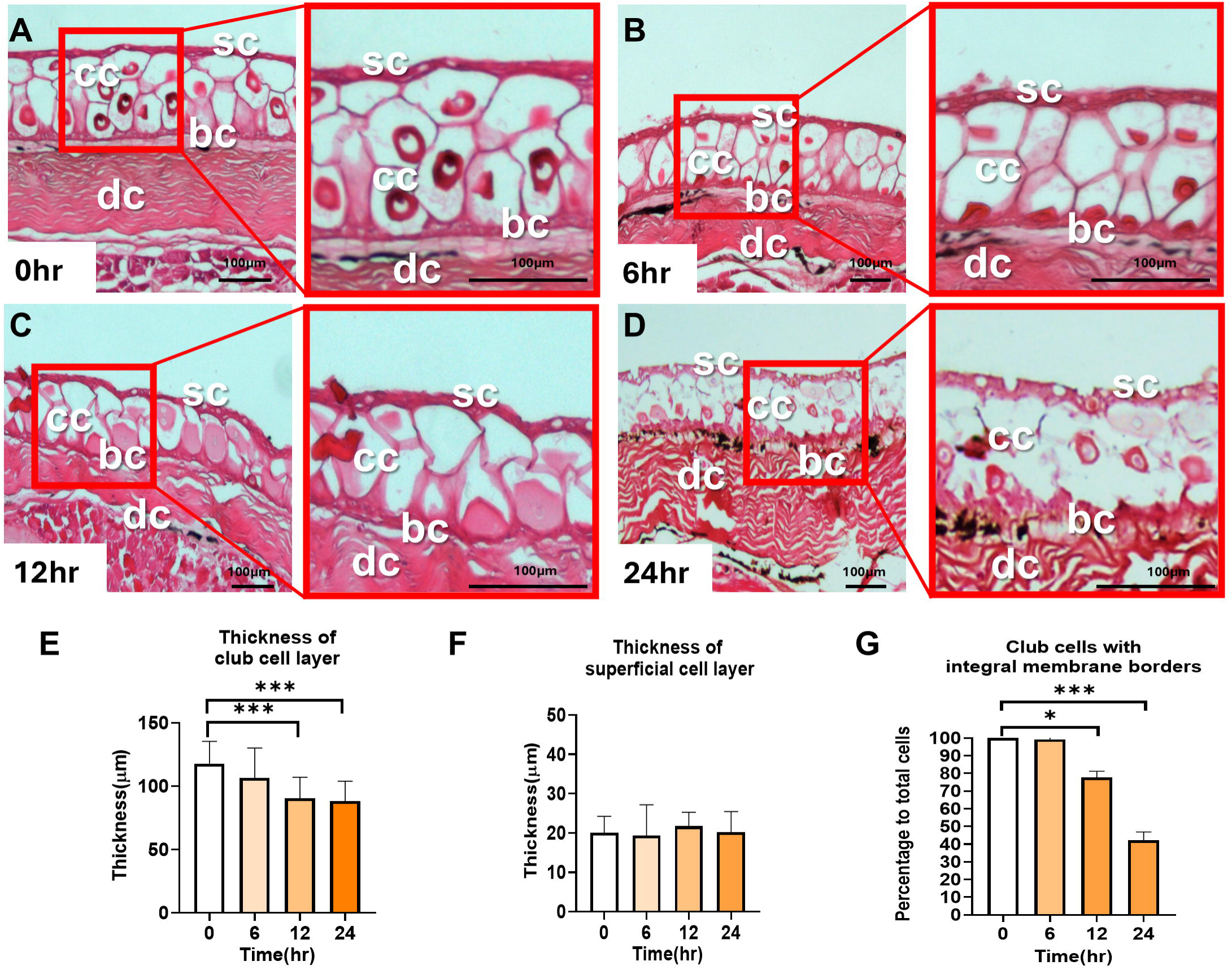
Evaluation of skin epithelial integrity over time under ammonia stress. Fresh skin was paraffin-preserved and sectioned. Tissue sections of ammonia-challenged and nonchallenged skin were stained with hematoxylin and eosin (H&E). Sections showing the superficial epithelial cells (sc), club cells (cc), basal epithelial cells (bc), and dense collagenous tissue (dc). (A–D) Low to high-magnification images of skin. Shown are representative images, Bar = 100 μm. Images are used to determine (E) the thickness of the club cell layer, (F) the thickness of the superficial cell layer, and (G) the percentage of club cells with integral membrane borders. Data are presented as mean ± SD. 3 fish with 2 tissue replicates from each were used. Morphometric evaluation was performed on 18 randomly selected fields per sample for each condition. All statistical significance was determined by one-way ANOVA followed by Tukey’s multiple comparisons test at * (p < 0.05) ** (p < 0.01) *** (p < 0.001).

### 3.4. Ammonia accumulates and decreases in striped catfish skin tissue

Ammonia accumulates in tissues correlates with exposure time [50, 51]. Thus, time–dependent accumulation of damage to tissue integrity can occur [52]. Since skin tissue showed subtle damage without loss of tissue integrity throughout the challenge, we hypothesized that ammonia does not accumulate or can be regulated to a non-damaging level. To evaluate the ammonia concentration in the skin tissue, Paraffin-sectioned skin tissue was stained by Nessler’s staining and imaged by a light microscope. We found that staining in skin tissue, particularly in the epidermis, increased after challenge, suggesting ammonia accumulation in the skin (Fig. 4A-D). We then examined the level and distribution of ammonia in the epidermis by color intensity using lookup tables (LUTs) in ImageJ. We found the superficial layer, but not the culb cell layer, accumulated the most ammonia. Surprisingly, although ammonia accumulated in the superficial cell layer from 0 to 12 hr, it was reduced and distributed more evenly from 12 to 24 hr (Fig. 4E-H). To confirm this observation, the linear-distance LUT color intensity values were quantified from the top to the bottom boundary of the superficial cell layer. The highest and lowest thresholds representing the least and most concentrated ammonia across the superficial layer from top to bottom were determined at 0 hr and among the challenge groups, respectively. Consistent with our observation, we found that staining reached its highest threshold corresponding to the most concentrated ammonia signal in the superficial layer and decreased towards the lowest threshold after 12 hr (Fig. 4I–L). Notably, we found that ammonia was most diluted at the outer edge of the superficial layer. These results showed that ammonia accumulated, followed by recovery in the skin during the acute ammonia stress. Direct measurement of skin tissue ammonia using an ammonia/ammonium assay kit (Bioworld Technology, China) further revealed that the skin ammonia concentration stopped increasing at 6 hr (Fig. S3), supporting the observation from Nessler’s staining. Taken together, ammonia in skin does not follow a time-dependent accumulation pattern but decreases during acute ammonia stress, thus suggesting an ammonia regulatory system in fish skin.

**Figure 4.**
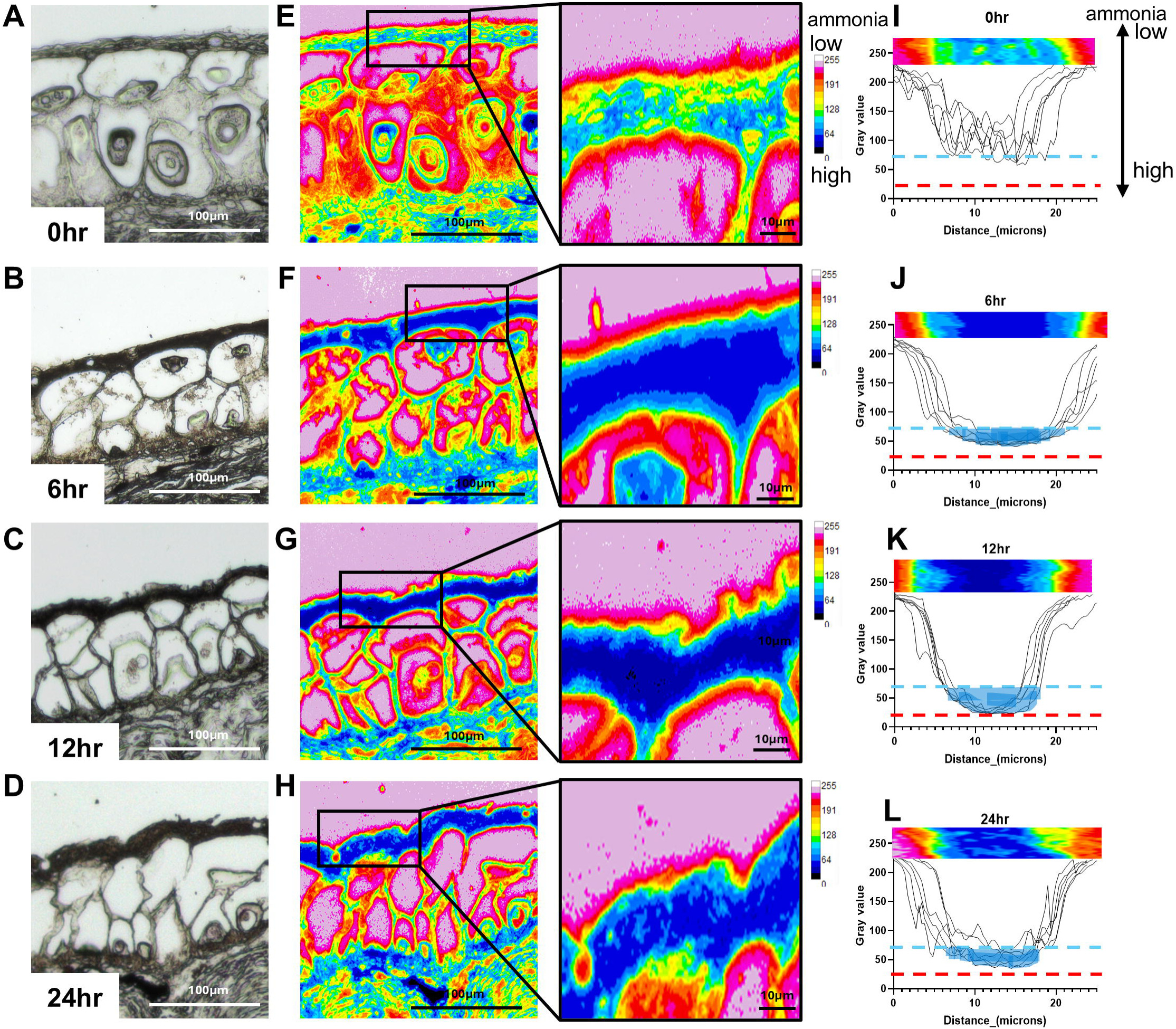
Evaluation of skin epithelial ammonia content over time under ammonia stress. Fresh skin was paraffin-preserved and sectioned. Tissue sections of ammonia-challenged and nonchallenged skin were stained with Nessler‘s reagent. (A-D) Images of Nesslar’s stain epithelial cells. Shown are representative images, Bar = 100 μm. (E-H) Images of LUT in epithelial cells, and magnification of superficial cells, were used to determine (I-L) the color intensity values of the linear distance in superficial cells. Bar = 100 μm, and 10 μm. 3 fish with 2 tissue replicates from each were used. The value was determined from 6 randomly selected fields in each condition.

### 3.5.1 Transcriptomic analysis of striped catfish skin under acute ammonia stress

Since references on the regulation of fish skin by ammonia are lacking, we sought to explore the hypothesized regulatory response using RNA-seq and transcriptomic profiling in fish skin. This approach allowed us to extensively scrutinize gene expression patterns that could show a molecular function for the regulatory pathway. RNA-seq reads from each time point were mapped to the striped catfish reference genome. Each sample yielded 23.7–45.78 million raw reads, and 18.20–39.39 million clean reads after trimming (Table S1). Clean read quality ranged from 97.10% to 98.85% for Q20 and 92.88% to 96.42% for Q30. GC content ranged from 48.23% to 53.81% (Table S2). Over 84% of reads were uniquely mapped (Table S1). Gene expression levels were calculated as counts per million (cpm) (Table S3).

#### 3.5.2 Skin transcriptomic profiles expand over ammonia exposure time

To analyze gene expression relations among samples, principal components analysis (PCA) was used to assess sample relationships. As no significant differences were observed among the non-challenged controls at each time point (Fig. S4), these were combined as a merged non-challenge control (mct) group for further analysis. We found that challenge groups showed greater variation over time than the mct group (Fig. 5A). Differential expression genes (DEGs) between the challenge and mct groups were identified using over-representation analysis (ORA) and visualized in volcano plots. At 6 hr, 267 DEGs were detected (41 down-regulated, 226 up-regulated; Fig. 5B). At 12 hr, 2670 DEGs were found (1190 down-regulated, 1480 up-regulated; Fig. 5C). At 24 hr, 8843 DEGs were identified (4508 down-regulated, 4335 up-regulated; Fig. 5D). Taken together, both the expression levels and number of DEGs increased over time, resulting in expansion of skin transcriptomic profiles over ammonia exposure time.

**Figure 5.**
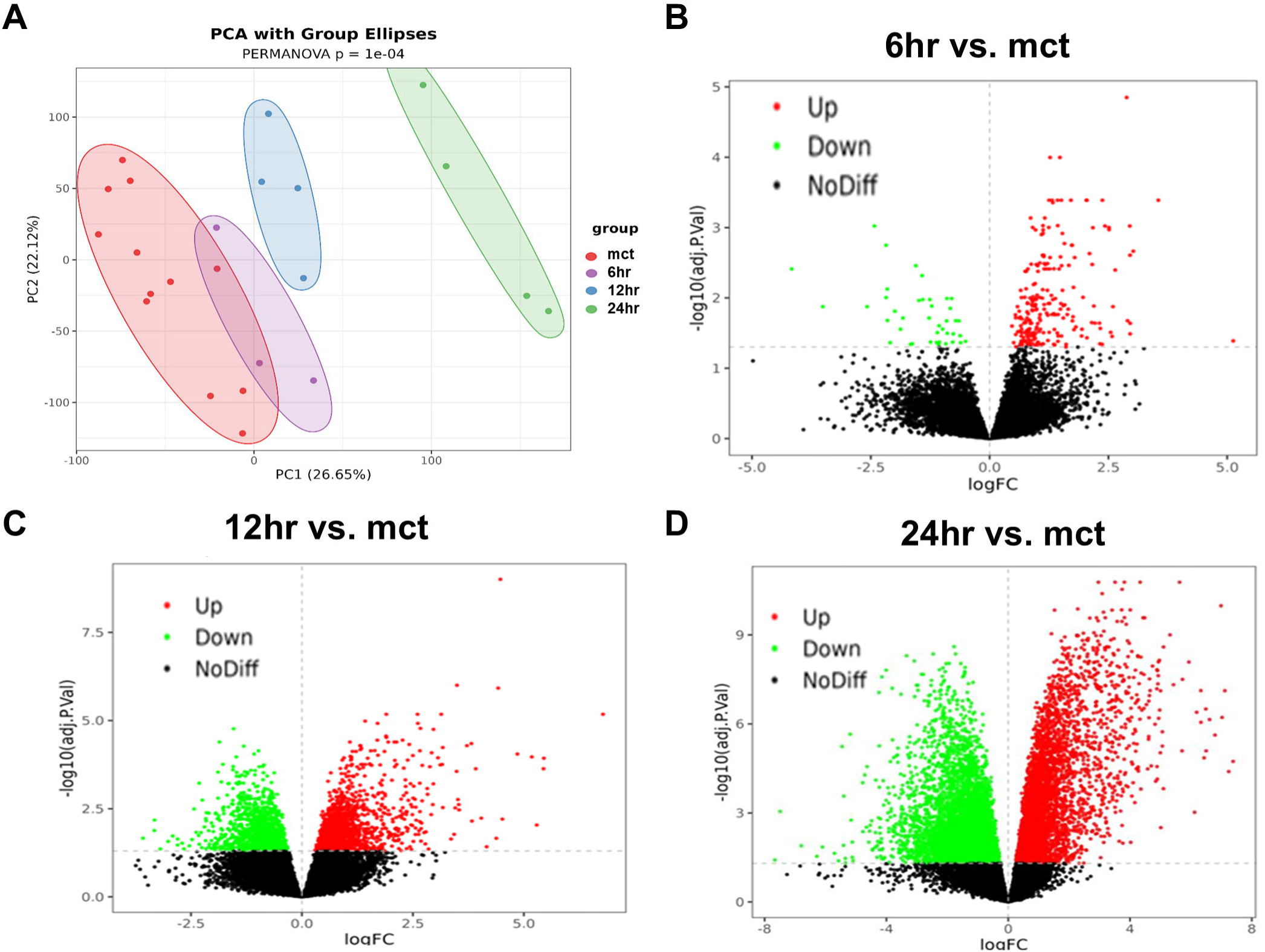
Analysis of striped catfish skin transcriptome difference over time under ammonia stress. Skin RNA from ammonia-challenged and non-challenged fish was extracted and subjected to RNA-seq. Fish skin (A) gene expression in PCA analysis. (B-D) Volcano plots illustrating log (FDR) about the log_2_(Fold change) for 6, 12, and 24 hr. Genes that passed the significance threshold FDR adjusted *P* < 0.05, and the expression cut-off | log_2_ (fold change) | ≥ 1 are colored red and green, while genes outside this range are colored black. N = 3 fish in each group.

### 3.6. Functional findings potentiate a distinct and immune-independent regulation

To further investigate the orchestration between expression levels and the number of DEGs, GO and KEGG enrichment analyses were performed using the Gene Ontology Consortium and the KEGG database [42–45]. Gene Set Enrichment Analysis (GSEA) was applied at each time point to compare Cellular Components (CC), Biological Processes (BP), and KEGG pathways between ammonia-challenged and mct groups. Firstly, the top 30 significantly enriched GO terms and KEGG pathways were visualized as heatmaps based on significant expression level (Fig. 6A–C). Significance levels across all three categories were unevenly distributed over time, suggesting that transcriptomic function levels do not follow a strictly time-dependent pattern. Sequentially, to examine the gene numbers, genes in each category were separated into up- and down-regulated groups. Each term at every time point was categorized into broader functional groups within existing categories. Based on shared parent terms in the GO and KEGG databases, CC terms were classified into mitochondria, organelle, cell membrane, nucleolus, and cell junction. BP terms were classified into metabolism, response to stimulus, cellular process, immune, and cell junction. KEGG terms were classified into metabolism, cellular process, genetic information processing, and immune groups. The numbers of up- and down-regulated genes in each category were summed and displayed as bar charts to examine the distribution of DEGs. The ratio of up-to-down-regulated gene count was also calculated to assess the regulation module over time (Fig. 6D–F). Similar to a heat map, we found that distributions and ratios across all three categories fluctuated over time, without a progressive trend over the 24 hr period. This data showed that a cellular function was consistently and significantly regulated by multiple genes, suggesting that regulation can likely be identified in function-based analysis. Moreover, the trend of the ratio over 24 hr can be classified into two categories. First, functional groups that were relatively unchanged within 24 hr, such as the cell junction in CC (Fig. 6D), the adhesion in KEGG (Fig. 6F), and the immune response in BP and KEGG (Fig. 6D and F). Second, functional groups with up-to-down-regulated gene count ratio peaked at 12 hr, such as mitochondria, organelle, and cell membrane in CC (Fig. 6D), metabolism in BP (Fig. 6E), and metabolism and genetic information processing in KEGG (Fig. 6F). Notably, the gene number and ratio change in CC were the most significant, whereas insignificant or unchanged in immune response. Hence, these data suggest a distinct cellular-component-based, immune-response-independent ammonia-regulation system in skin within 24 hr.

**Figure 6.**
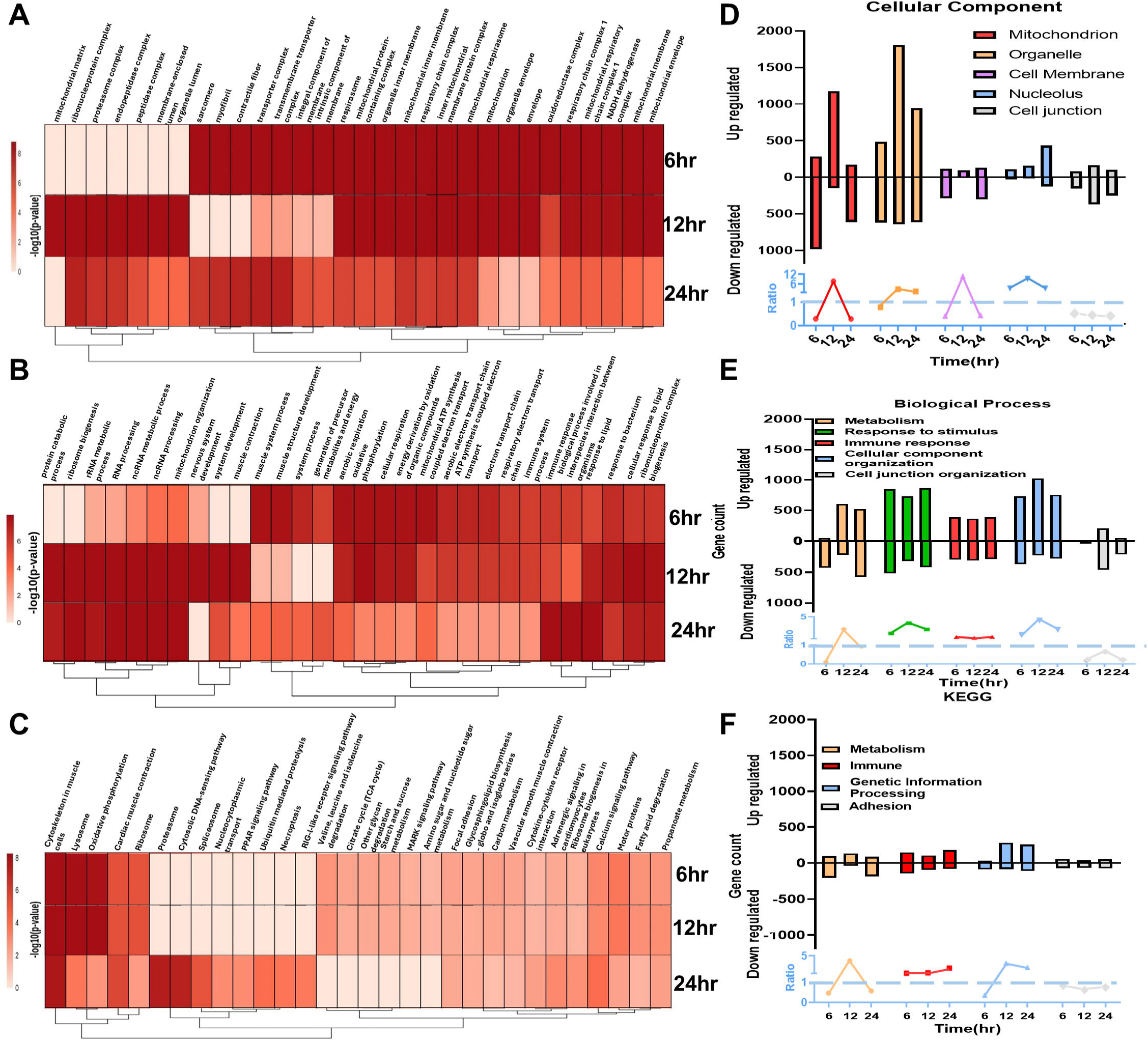
Analysis of the gene function of striped catfish skin transcriptome over time under ammonia stress. (A) Heatmap and (B) bar chart of 5 categories of the top 30 enriched cellular components. (C) Heatmap and (D) bar chart of 5 categories of the top 30 enriched biological processes. (E) Heatmap and (F) bar chart of 4 categories of the top 30 enriched KEGG. Bar chart of fish skin DEGs was first separated into down-regulated genes (down portions of the y-axis) and up-regulated genes (up portions of the y-axis). N = 3 fish in each group.

### 3.7. Increased small vesicles in superficial cells in striped catfish skin under acute ammonia challenge

Since RNA-seq indicated that cellular components exhibit the promised variation and Nesslar’s staining suggested that the superficial layer could regulate ammonia concentration, we further explored the cellular components of superficial cells to investigate this regulation. Transmission Electron Microscopy (TEM) was used to examine cell components in detail. In line with the paraffin-based findings, we found that the superficial integrity remained unchanged. However, we found that increased vacuoles and vesicles in cells at the outermost superficial layer from 6 to 24 hr compared to cells at 0 hr (Fig. 7A). The vacuole size (0.1-0.4 µm^2^) was found comparable to that of mitochondria observed at 0 hr. Furthermore, we found mitochondria gradually lost integrity and degraded as the challenge progressed (Fig. 7B), the data implied that mitochondrial damage and/or recycling may be associated with increasing vacuole number in the skin under ammonia stress. The vesicle size (0.02-0.03 µm^2^) was found to be comparable to that of reported endosome/exosome vesicles [53]. Notably, we found these vesicles not only significantly increased but also distributed towards the surface of outermost epithelial cells within the superficial layer (Fig. 7C). With the number of vacuoles and vesicles being quantified, we found that both vacuole and vesicle number significantly increased at 12 hr and remained similar at 24 hr (Fig. 7D & E). Consistently, negative correlation between the number of vacuoles and mitochondira were found in a time-dependent trend (Fig. 7F), supporting vacuoles’ origin from mitochodia degredation. Since vesicles are known for their ability to transport molecules, we explored this possibility by evaluating exosome- and endosome-related gene expression from the transcriptomic profile. We found that the expression of both types of vesicles increased at various levels. However, the number and level of exosome-related but not endosome-related genes increased more significantly within 24 hr (Table 3), indicating that the observed vesicles may primarily be exosome-based. Together, these data support the hypothesis that exosome vesicles may regulate skin tissue ammonia accumulation by transporting ammonia out of the cell.

**Figure 7.**
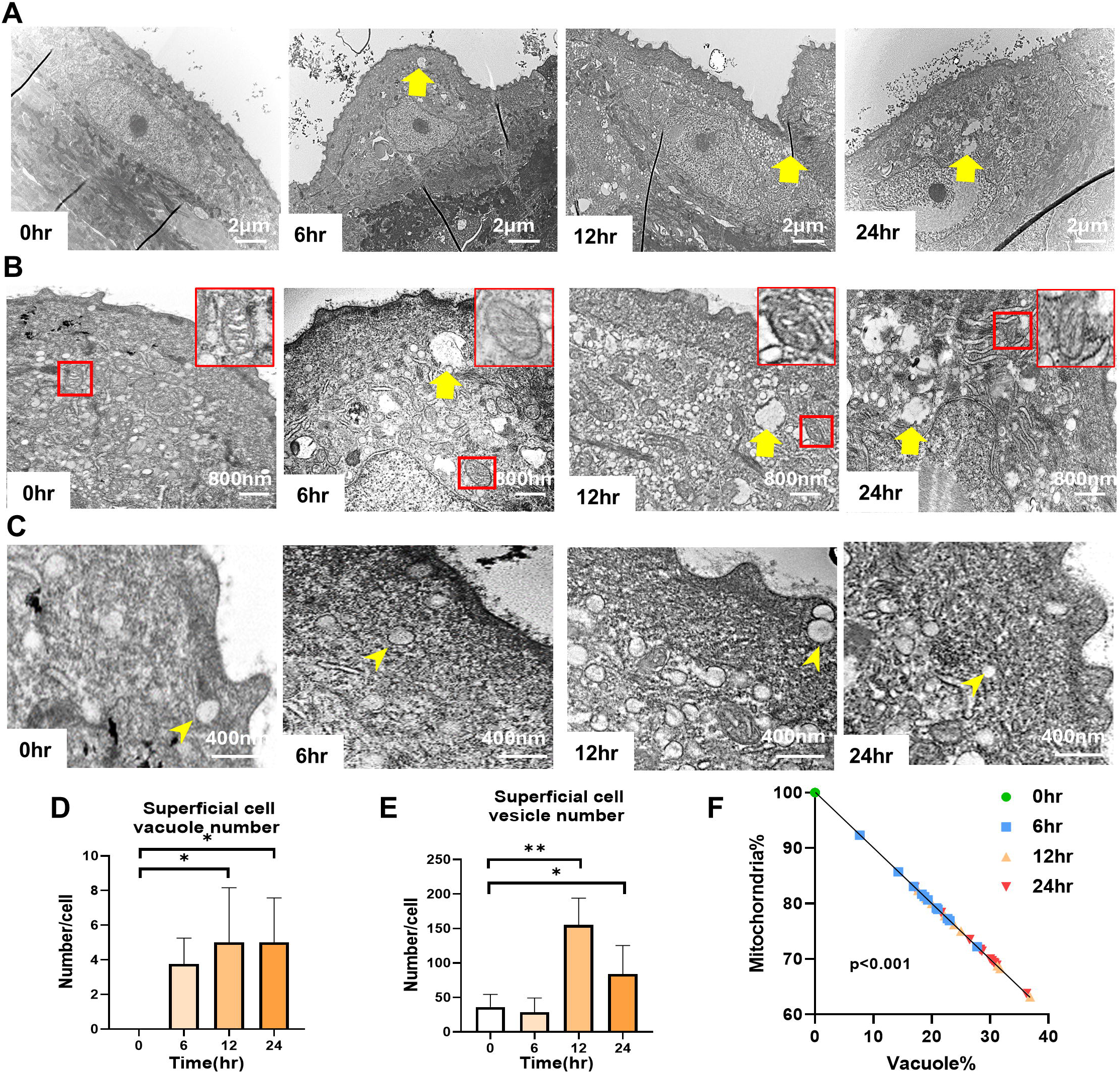
Exploration of skin integrity and skin structure over time under ammonia stress. Transmission electron microscopy (TEM) images of fresh skin were taken. Sections showing the vacuole 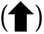 and vesicle 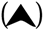. Images of (A) the skin superficial cell integrity, Bar □= 2 μm, (B-C) the upper layer of the superficial cells, bar =800 nm and 400 nm. Images are used to determine the (D) vacuole and the (E) vesicle number, and (F) relative proportion of mitochondria and vacuole in superficial cells. Data are presented as mean ± SD. 3 fish with 2 tissue replicates from each were used. Morphometric evaluation was performed on 12 randomly selected fields per sample for each condition. All statistical significance was determined by one-way ANOVA followed by Tukey’s multiple comparisons test at * (p < 0.05) ** (p < 0.01).

**Table 3.**
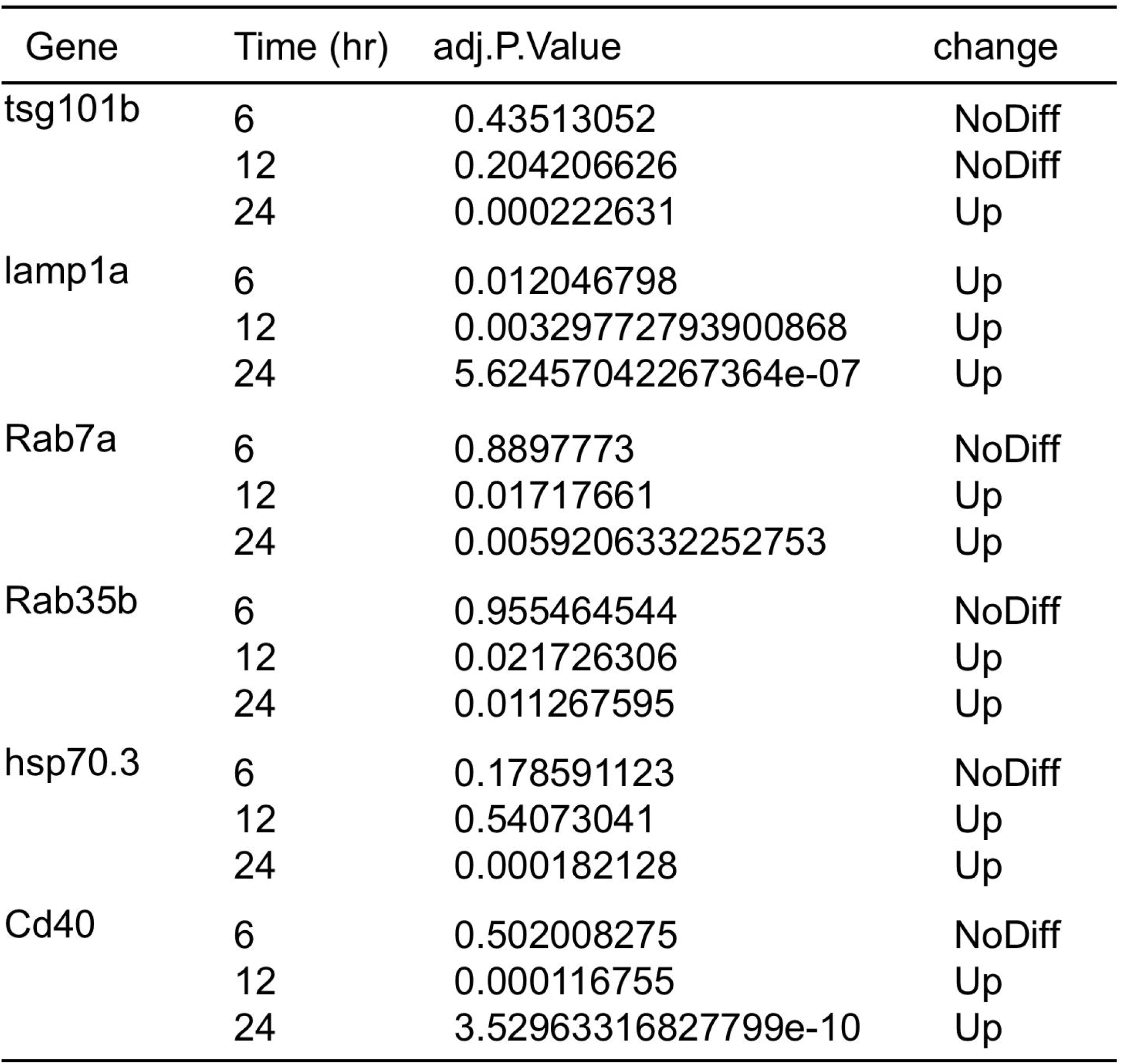
Exosome-related gene expression.

| Gene | Time (hr) | adj.P.Value | change |
| --- | --- | --- | --- |
| tsg101b | 6 | 0.43513052 | NoDiff |
|  | 12 | 0.204206626 | NoDiff |
|  | 24 | 0.000222631 | Up |
| lamp1a | 6 | 0.012046798 | Up |
|  | 12 | 0.00329772793900868 | Up |
|  | 24 | 5.62457042267364e-07 | Up |
| Rab7a | 6 | 0.8897773 | NoDiff |
|  | 12 | 0.01717661 | Up |
|  | 24 | 0.0059206332252753 | Up |
| Rab35b | 6 | 0.955464544 | NoDiff |
|  | 12 | 0.021726306 | Up |
|  | 24 | 0.011267595 | Up |
| hsp70.3 | 6 | 0.178591123 | NoDiff |
|  | 12 | 0.54073041 | Up |
|  | 24 | 0.000182128 | Up |
| Cd40 | 6 | 0.502008275 | NoDiff |
|  | 12 | 0.000116755 | Up |
|  | 24 | 3.52963316827799e-10 | Up |

**Table 4.** Endosome-related gene.

| Gene | Time (hr) | adj.P.Val | change |
| --- | --- | --- | --- |
| eea1 | 6 | 0.745511441 | NoDiff |
|  | 12 | 0.494519717 | NoDiff |
|  | 24 | 0.009836776 | Up |
| vps33b | 6 | 0.975844838 | NoDiff |
|  | 12 | 0.910189217 | NoDiff |
|  | 24 | 0.057596406 | NoDiff |
| entr1 | 6 | 0.758951084 | NoDiff |
|  | 12 | 0.65590799 | NoDiff |
|  | 24 | 0.98555433 | NoDiff |
| elapor1 | 6 | 0.669846754 | NoDiff |
|  | 12 | 0.930104819 | NoDiff |
|  | 24 | 0.243976592 | NoDiff |
| elapor2a | 6 | 0.256401363 | NoDiff |
|  | 12 | 0.25148365 | NoDiff |
|  | 24 | 0.947773082 | NoDiff |
| mon2 | 6 | 0.85923077 | NoDiff |
|  | 12 | 0.063277122 | NoDiff |
|  | 24 | 0.44136925 | NoDiff |

### 3.8. Exosomes in skin epithelial cells play a role in regulating ammonia excretion

To verify whether ammonia concentration was regulated via exosomes in skin epithelia, an established striped catfish skin epithelial cell line was used [47] with or without exosome inhibitor GW4869 under acute ammonia stress. Basically, cells were pretreated with 1 hr of GW4869 and then subjected to a 6 hr challenge with 40 mg/L ammonia, followed by a 1 hr recovery period with ammonia-free media (Fig. 8A). The ammonia concentration in both media and cells was measured using Nitrogen-Ammonia Reagent Set (Hach, USA) to determine the exosomal inhibition effect of GW4869 on ammonia. We first examined whether cell line viability was affected by ammonia and/or GW4869. We found no significant change in cell morphology in any group under the light microscope (Fig. S5), and no significant cell exfoliation was detected (Fig. 8B & C), confirming the validity of the experimental results obtained with ammonia and GW4869. To further verify that GW4869 did not affect ammonia measurement, we compared supernatant and intracellular ammonia concentrations in cells treated with or without GW4869 under conditions without ammonia stress. No significant differences were observed after 6 hr of challenge and 1 hr of recovery (Fig. 8D–G), confirming the applicability of GW4869. Subsequently, cells exposed to ammonia with or without GW4869 were examined. We found that GW4869 treatment significantly reduced the ammonia levels in the supernatant to ∼3 mg/L after 6 hr, concurrently leading to an accumulation of ∼1.6 mg/L intracellular ammonia (Fig. 8D & E). A similar effect was observed after 1 hr of recovery (Fig. 8F & G). These findings demonstrate that inhibition of exosome activity significantly reduces ammonia excretion from skin epithelial cells, thus suggesting that exosomes play a role in regulating ammonia excretion in striped catfish skin.

**Figure 8.**
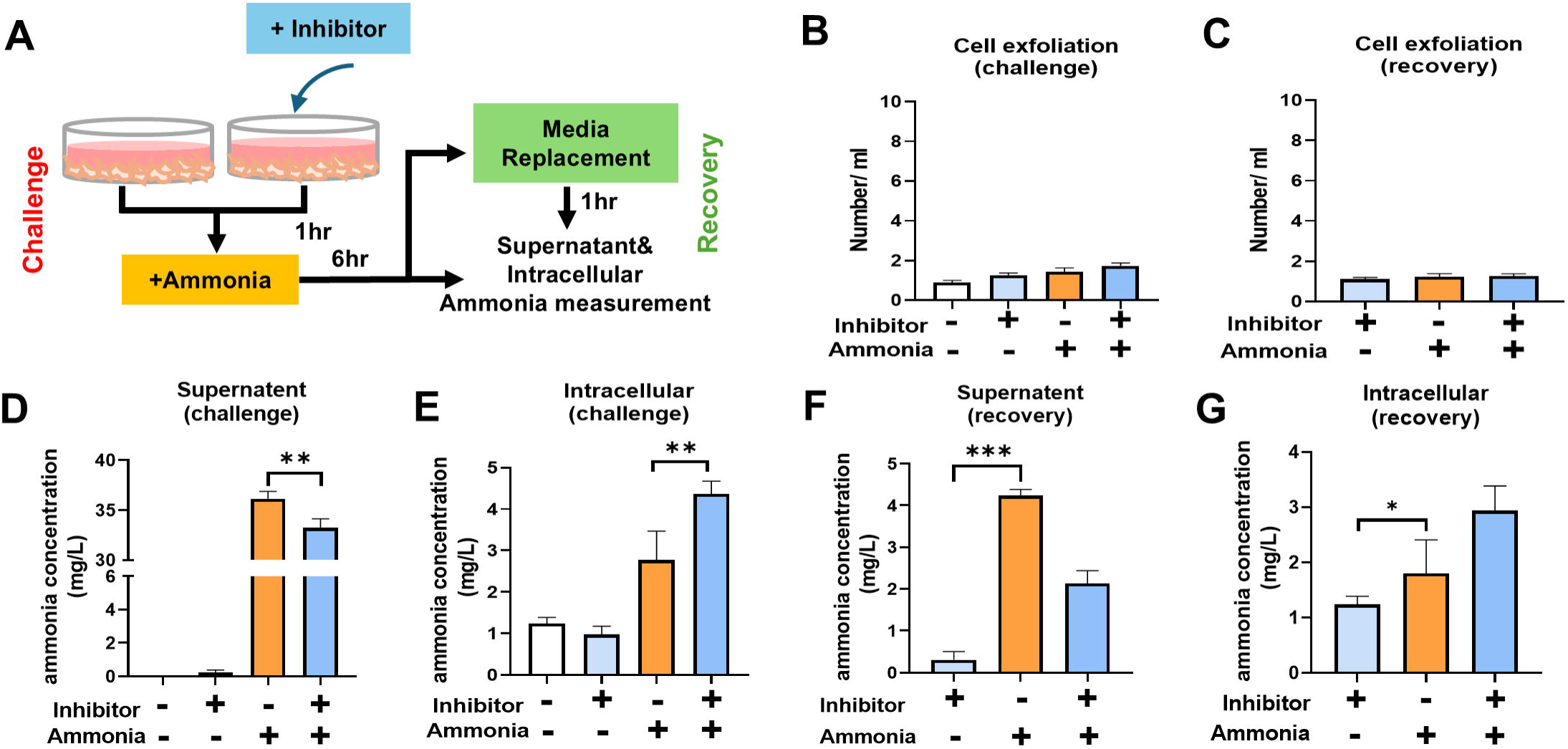
Validation of skin exosome-associated ammonia excretion. The striped catfish skin epithelial cell line with inhibited exosome release was challenged with ammonia for 6 hr, followed by a 1 hr recovery period. (A) Illustration of the experiment. (B–C) Quantification of exfoliated cells after 6 hr challenge and 1 hr recovery. Ammonia concentrations in (D) supernatant and (E) cells after 6 hr challenge, and in (F) supernatant and (G) cells after 1hr recovery. Data are presented as mean ± SD. N = 3 in each group. All statistical significance was determined by one-way ANOVA followed by Tukey’s multiple comparisons test at * (p < 0.05), ** (p < 0.01), *** (p < 0.001).

## 4. Discussion

The present study investigates the skin responses of striped catfish to acute ammonia stress. The findings demonstrate that the skin, though inflamed, maintains tissue integrity, potentially attributed to an unexplored exosome-based ammonia excretion system. While most studies reported only undamaged or undefined skin observations, our study provides mechanistic insights into how fish skin remains mostly undamaged under ammonia stress. To the best of our knowledge, this is the first study to highlight striped catfish’s skin responses and their regulation of ammonia, providing valuable information for future aquaculture management.

Ammonia can induce alterations in fish physiology and immunology that can impact tissue histology [13]. In physiological studies, Wicks et. al found swimming speed decreases in coho salmon *O. kisutch* under 0.08 mg/L ammonia stress for 24 hr [54]. A more severe decrease in general movement was found in red goldfish, ranging from 26.2 mg/L of ammonia stress for 24 hr [55]. In striped catfish, we also observed decreased physical activity under 40 mg/L TAN stress, indicating the similarity of general damage from ammonia. In immunological findings, striped catfish liver, spleen, and skin all showed significant inflammation under acute ammonia stress. The liver and spleen are two internal organs most susceptible to ammonia stress. Yellowfin tuna *T. albacares,* and Nile tilapia *O. niloticus* showed more than 3- and 2-fold change in liver inflammation under even 5 mg/L ammonia stress for 24 hr [56, 57]. Moreover, 5 mg/L ammonia stress for 24 hr can induce more than 20-fold increase in splenic inflammation in Yellowfin tuna *T. albacares* [58, 59]. Consistent with our study, these suggest that the Striped catfish’s overall health response to ammonia is similar to that of other fish. Particularly, the skin of striped catfish, though with increased inflammation signals, remains structurally unaffected under acute ammonia stress, different from increased skin intercellular spaces and filament cell vacuolization, which have been reported in Gilhead bream *S. aurata* under 3 to 12 mg/L TAN stress for 24 hr [28]. However, roughened apical epithelial surfaces and cytoplasmic vacuolation of the mitochondria found in skin epithelial cells of Japanese bream *N. japonica* larvae under 0.26 mg/L TAN stress resemble our findings [60]. These indicate that teleost fish skin has responses to ammonia at different levels, and striped catfish skin under acute ammonia stress maintains the structurally different but cellularly comparable to other teleost fish.

Notably, we found that the club cell layer in the skin dermis showed cell membrane thinning or rupture. Club cells have been reported to release their contents after their cell membrane is incomplete and ruptured under stress [61, 62]. The content forms a protective layer over the deeper epidermal layers and helps seal damage [63]. For instance, in the Air-breathing catfish *C. batrachus*, skin club cells can confluence with neighboring cells under detergent stress [64]. Therefore, club cell thinning or rupture in striped catfish skin may pose a protective mechanism rather than damage under acute ammonia stress. However, how the skin copes with ammonia and/or adapts for a longer period is unknown. Further investigation into the long-term effects of ammonia exposure in striped catfish may provide the answer to how skin adaptively survive in ammonia stress.

Rhesus (Rh) glycoproteins play a crucial role in the ammonia excretion process [65]. These proteins are typically distributed in the cell membrane and mitochondrial membrane of epithelial cells [66, 67], where they facilitate the transport of ammonia from the blood and cytoplasm to the external environment. Previous studies have reported upregulation of Rh glycoproteins expression in the skin of Mangrove killifish *K. marmoratus* under 28 mg/L TAN for 5 days and Rainbow trout *O. mykiss* 21 mg/L TAN for 12 and 48 hr [66, 68, 69]. In Zebra fish *D. rerio,* skin was also found to excrete ammonia via Rh glycoproteins under normal conditions [68]. We also observed several Rh-related genes with an increasing trend, though not significant at 6 and 12 hr (Table S4), leaving two possibilities: Rh glycoproteins are not the dominant pathway for ammonia excretion in striped catfish skin, or they require a longer exposure time to exert a significant effect. In our study, an inflamed skin with an increase in cellular vesicles migrating toward the superficial cell membrane in epidermis (Fig. 7 & S4) addressed the first possibility. Supported by a significant increase in exosome-related gene expression (Table S4) and confirming exosomes’ role involved in ammonia excretion in striped catfish skin (Fig. 9), we verified that an additional ammonia excretion system in skin besides Rh glycoproteins. Based on studies showing ammonia disturbance in the cytoplasm, we propose that ammonia may be engulfed into exosomes during their formation. Two potential sources of these exosomes: one originating from late endosomes and endoplasmic reticulum (ER), and the other possibly derived from the mitochondrial membranes [70–72]. Previous studies have demonstrated that exosomal cargo can be transferred from the ER and cytoplasm to late endosomes, where it then becomes exosomes [73]; therefore, we hypothesize that ammonia could be incorporated via a similar pathway. Moreover, mitochondria are known to be associated with Rhesus (Rh) glycoproteins and can accumulate ammonia. Damaged mitochondria and mitochondrial components can generate autophagic vesicles, some of which may fuse with late endosomes, suggesting that mitochondrial substances—including ammonia—could be packaged within exosomes and subsequently secreted [74, 75]. It would be interesting and also essential to investigate how ammonia is transported into vesicles and exosomes and then secreted into the extracellular space to complete the pathway in this exosome-based ammonia secretion system.

Even though ammonia could be excreted out of the skin, the maintenance of skin survival and metabolism from ammonia impact is required. Inflammation can help mitigate ammonia-induced damage, as reported in immune-related organs such as the liver, spleen, kidneys, and gills in various fish species under acute ammonia stress [76–78]. In striped catfish, skin, liver, and spleen were observed with comparable inflammatory responses (Fig. S2). In addition, inflammation and oxidative stress may trigger autophagy as a compensatory mechanism to prevent further cell death [79–81]. Ammonia-induced autophagy was reported and shown to protect Yellow catfish *P. fulvidraco* from liver inflammation under a half-lethal dose of 125 mg/L TAN stress for 48 hr [25]. Our study found potential mitophagy formation, mitochondria-based autophagy, under inflamed skin by TEM (Fig. S6). Mitophagy is also known to maintain cellular homeostasis and prevent apoptosis [82]. If so, these mitophagy vacuoles likely represent a regulated cellular response to acute ammonia stress. Together, the inflammation and mitophagy may be protective or supported by additional mechanisms that help maintain skin integrity. However, because our observations were limited to acute conditions, it remains unclear whether striped catfish skin would develop prolonged or harmful inflammation under extended ammonia stress. Other protective mechanisms—such as metabolic regulation—may also contribute to the skin’s response to acute ammonia stress.

Most fish can also detoxify ammonia in addition to excretion across the epithelium of skin and gill by converting ammonia into urea and glutamine [67]. Previous studies have shown that ammonia metabolism–related genes involved in amino acid metabolism—such as amino sugar and nucleotide sugar metabolism (including fructose and mannose metabolism and glycolysis), alanine, aspartate and glutamate metabolism, arginine and proline metabolism, and glutamine biosynthesis—are upregulated in the liver of Large-scale loach *P. dabryanus* under 420 mg/L TAN stress for 48 hr [20]. Similarly, Asian seabass *L. calcarifer* exhibited enhanced glutamate metabolism under 5.58 mg/L ammonia stress for 60 days [83]. These findings suggest that fish can mitigate ammonia toxicity by metabolizing ammonia. In striped catfish, we found that ammonia metabolism–associated pathways, including amino sugar and nucleotide sugar metabolism and glutamate metabolism linked to glutamine synthesis, were significantly upregulated between 6 and 12 hr (Fig. 5C & Table S4), indicating potential ammonia conversion as another ammonia detoxifying method in skin tissue. Yet, the long-term effects reported in previous studies still need to be clarified to confirm the detoxification mechanism in the skin. In addition to directly metabolizing ammonia, fish may also enhance glucose and protein metabolism to meet the elevated energy demand during ammonia stress. For instance, Yellow catfish *P. fulvidraco* showed increased serum glucose levels under 2.65 mg/L ammonia stress for 24 hr [84], while common Carp *C. carpio* exhibited a similar response under 0.823 mg/L TAN for only 3 hr [84, 85]. In striped catfish skin, genes associated with glucose metabolism were also elevated under acute ammonia stress for 24hr (Table S4), indicating energy elevation for maintenance of tissue or cell integrity. Together, striped catfish skin, besides exosomal vesicles as an ammonia excreting system, may also exert acute ammonia stress regulation on maintaining tissue and cell integrity through inflammation, ammonia metabolism, and increased metabolic rate. How the orchestration of these regulatory systems renders the skin tissue’s survival or even renewal is of importance in a future perspective.

Despite the new findings in this study, this study has some limitations that extensive research could address. First, lack of regulatory information in each layer of skin. Fish skin comprises both the epidermis and dermis, which differ in structure and function [29, 86]. These two layers mount distinct immune responses [31]. In our histological observations, we found that epithelial cell types—including club cells and superficial cells—exhibit distinct characteristics. Spatial transcriptomics can be applied to resolve cell-type-specific gene expression within intact skin tissue, as demonstrated in studies of Atlantic salmon *S. salar* [87]. Second, skin cell renewal under ammonia stress remains unclear. Tissue repair requires a continuous supply of new cells to replace damaged ones [88]. In teleost, inflammatory and immune responses initiate regeneration, a critical step in subsequent tissue repair [89]. Previous studies have shown that skin epithelial thickness and mucus cell numbers increase after 4 days of ammonia exposure, suggesting partial regeneration even before the complete replacement of old cells [35, 52]. In striped catfish skin, we observed immune and inflammatory responses, and the MAPK pathway—known to regulate differentiation, survival, proliferation, and development [90], were significantly upregulated, implying that the skin may have entered the early phase of cell renewal (Fig. 5). Although some fibroblast growth factor genes were upregulated, others were downregulated, and several keratin and epidermal growth factor–related genes also showed decreased expression (Table S4), implying that active cell renewal had not yet begun under acute ammonia stress in this study design focusing solely on acute responses. Third, whether the skin microbiome is involved in the regulation of skin responses is unknown. The skin is covered by mucus that resides with the skin microbiome [91]. The composition of the skin microbiome and the metabolites present in the mucus are closely associated with the health status of the host [92]. Consequently, the host skin and its microbiome likely engage in continuous and complex interactions, particularly during stress conditions [93]. Previous studies demonstrated that ammonia exposure for more than 3 days can alter the fish skin microbiome, leading to an increase in opportunistic pathogens and inducing host inflammation [94–96]. Therefore, host health resulting from microbiome dysbiosis cannot be ruled out [91]. In contrast, studies also show that the microbiome assists the host fish in the detoxification of ammonia [97]. Moreover, as most existing studies have focused on microbiome responses under chronic ammonia exposure, it would be of great interest to investigate how the skin microbiome changes under acute ammonia stress.

## 5. Conclusion

Taken together, our findings not only report the general responses of striped catfish to acute ammonia stress but also reveal a novel exosome-mediated mechanism by which fish skin excretes ammonia and maintains homeostasis. Integrated with immune and metabolic functions, this study provides new insights into fish physiological adaptation and potential strategies to mitigate ammonia toxicity in aquaculture.

## CRediT statement

Wei-Hsuan Hsiao: Data curation, Formal analysis, Investigation, Methodology, Validation, Visualization, Writing—original draft.

Shun-Yi Lin: Data analysis

Ying-Hsien Wu: Methodology

Han-Chen Ho: Methodology, Data curation, Writing—review & editing

Tsung-Lin Liu: Writing—review & editing

Liang-Chun Wang: Supervision, Writing—review & editing, Funding acquisition

## Funding

This research was supported by grant (114-2813-C-110 -022 -B) and (113-2313-B-110 -005 -MY3) from the National Science and Technology Council (NSTC), funded by Taiwan, R.O.C.

## Declaration of competing interest

The authors declare that they have no competing financial interests or personal relationships that could have influenced the work presented in this study.

## Acknowledgement

We thank the Institutional Animal Care and Use Committee (National Sun Yat-Sen University) for its kind assistance, and the Department of Marine Biotechnology and Resources (National Sun Yat-Sen University) for providing the fluorescent microscopy and aquaculture system.

## Notes

### Competing Interest Statement

The authors have declared no competing interest.

### Summary of Updates

Modify the image number associated with Results. adding the author name in author lists

