## supplemental figures for "Investigating the skin response of Striped Catfish to acute ammonia stress reveals a potential exosome-based ammonia excretion system"

### Slide 1
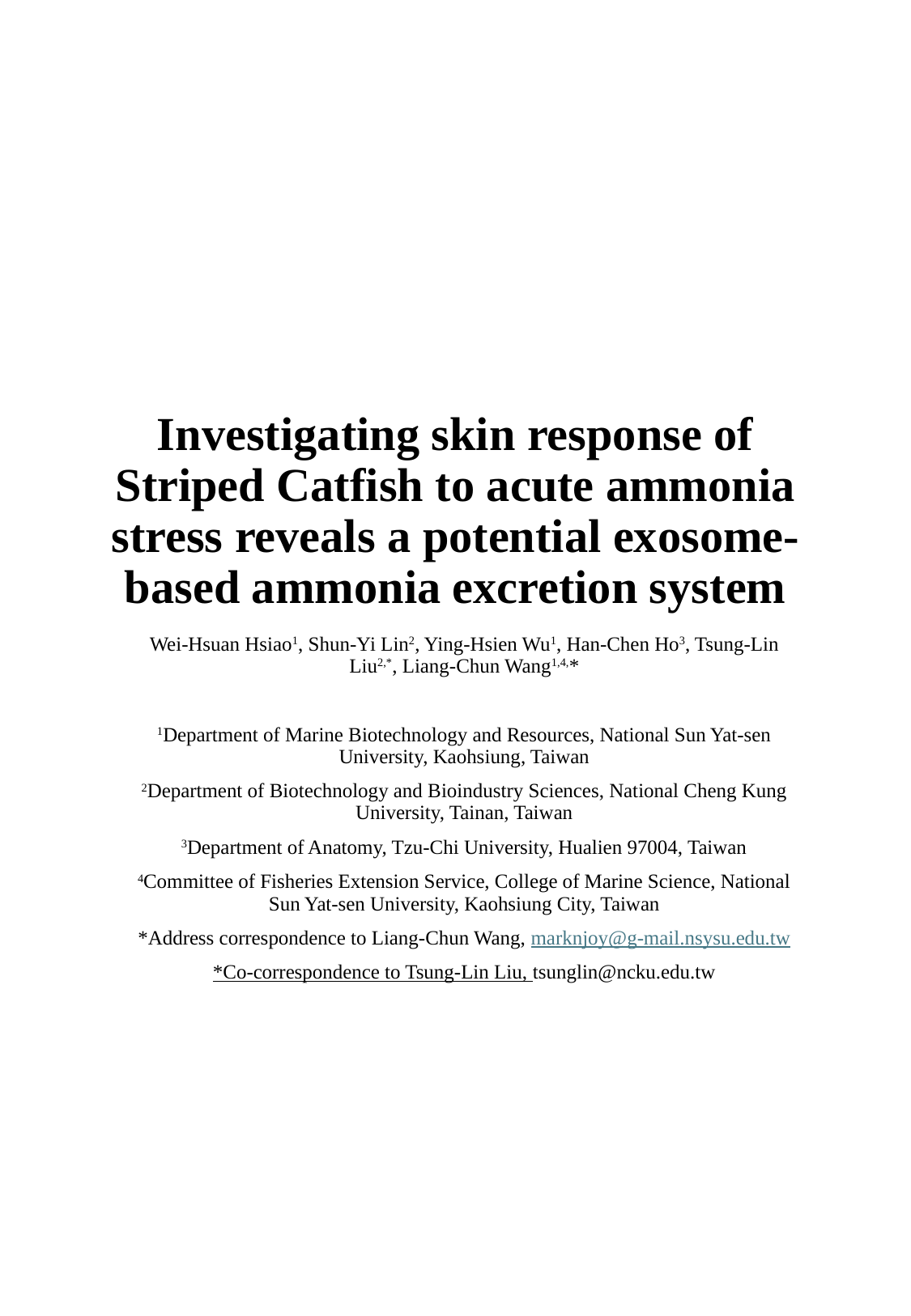

#### Investigating skin response of Striped Catfish to acute ammonia stress reveals a potential exosome-based ammonia excretion system
Wei-Hsuan Hsiao^1^, Shun-Yi Lin^2^, Ying-Hsien Wu^1^, Han-Chen Ho^3^, Tsung-Lin Liu^2,*^, Liang-Chun Wang^1,4,^*
^1^Department of Marine Biotechnology and Resources, National Sun Yat-sen University, Kaohsiung, Taiwan
^2^Department of Biotechnology and Bioindustry Sciences, National Cheng Kung University, Tainan, Taiwan
^3^Department of Anatomy, Tzu-Chi University, Hualien 97004, Taiwan
^4^Committee of Fisheries Extension Service, College of Marine Science, National Sun Yat-sen University, Kaohsiung City, Taiwan

### Slide 2
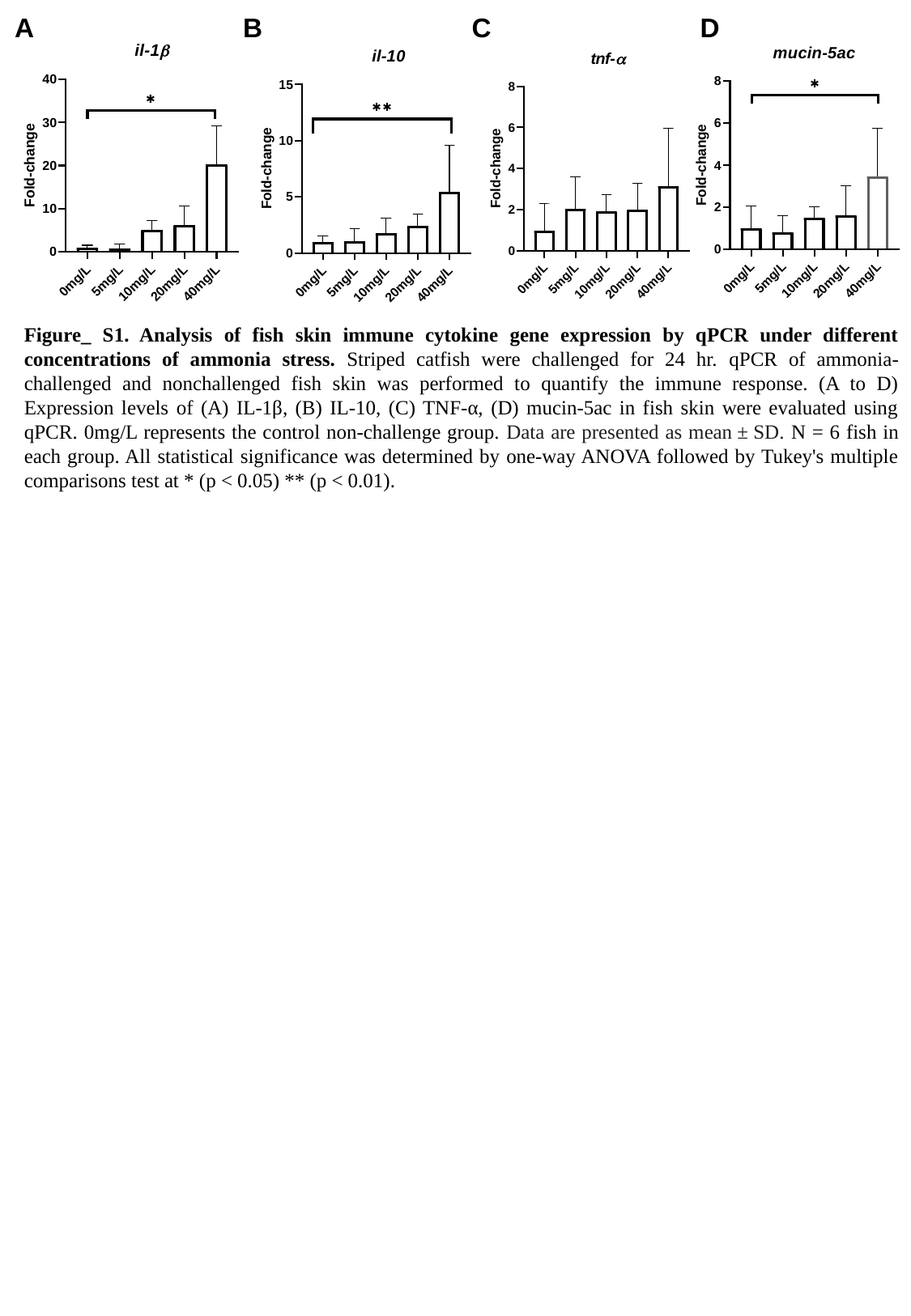

A
B
C
D
Figure_ S1. Analysis of fish skin immune cytokine gene expression by qPCR under different concentrations of ammonia stress. Striped catfish were challenged for 24 hr. qPCR of ammonia-challenged and nonchallenged fish skin was performed to quantify the immune response. (A to D) Expression levels of (A) IL-1β, (B) IL-10, (C) TNF-α, (D) mucin-5ac in fish skin were evaluated using qPCR. 0mg/L represents the control non-challenge group. Data are presented as mean ± SD. N = 6 fish in each group. All statistical significance was determined by one-way ANOVA followed by Tukey's multiple comparisons test at * (p < 0.05) ** (p < 0.01).

### Slide 3
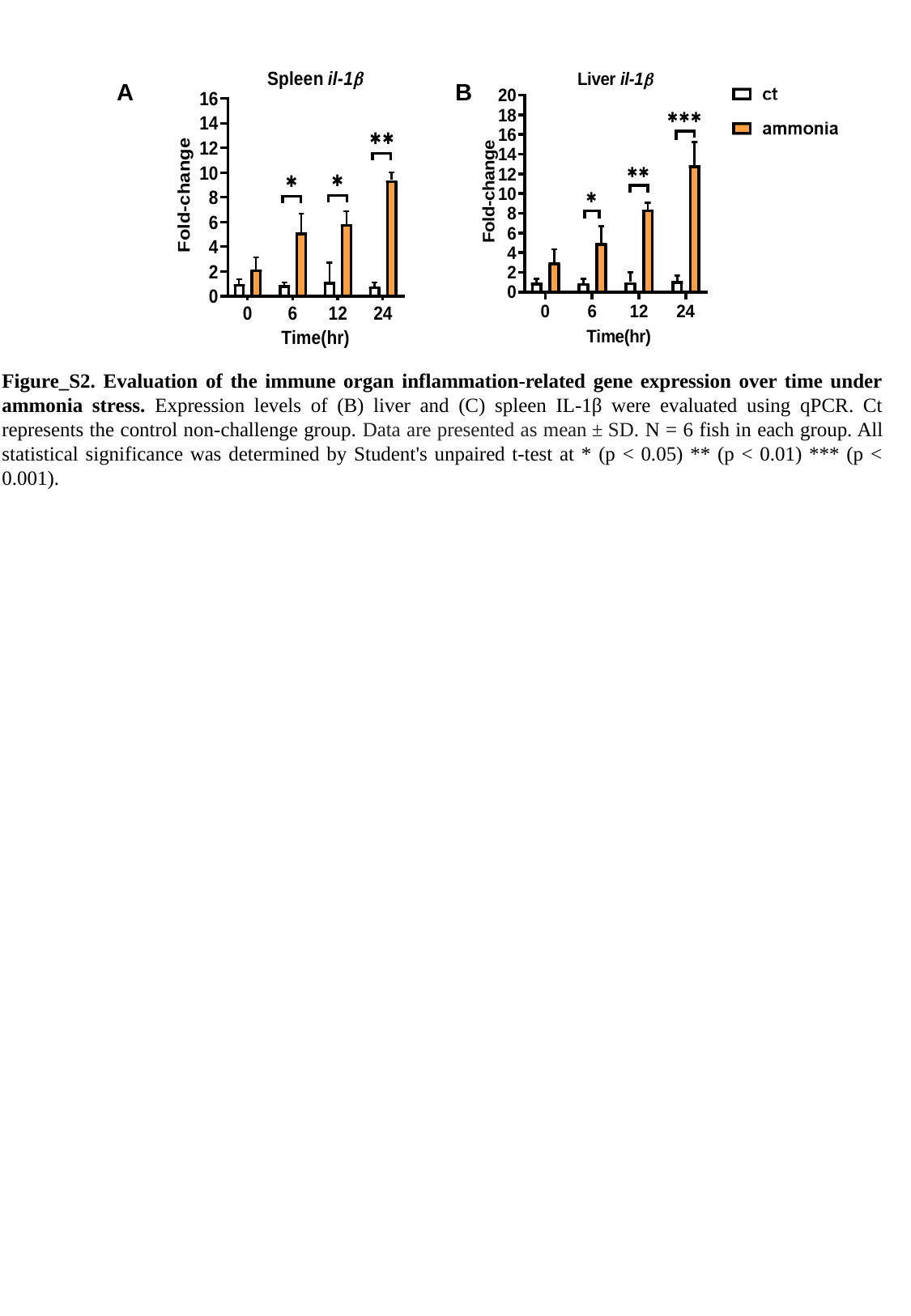

A
B
Figure_S2. Evaluation of the immune organ inflammation-related gene expression over time under ammonia stress. Expression levels of (B) liver and (C) spleen IL-1β were evaluated using qPCR. Ct represents the control non-challenge group. Data are presented as mean ± SD. N = 6 fish in each group. All statistical significance was determined by Student's unpaired t-test at * (p < 0.05) ** (p < 0.01) *** (p < 0.001).

### Slide 4
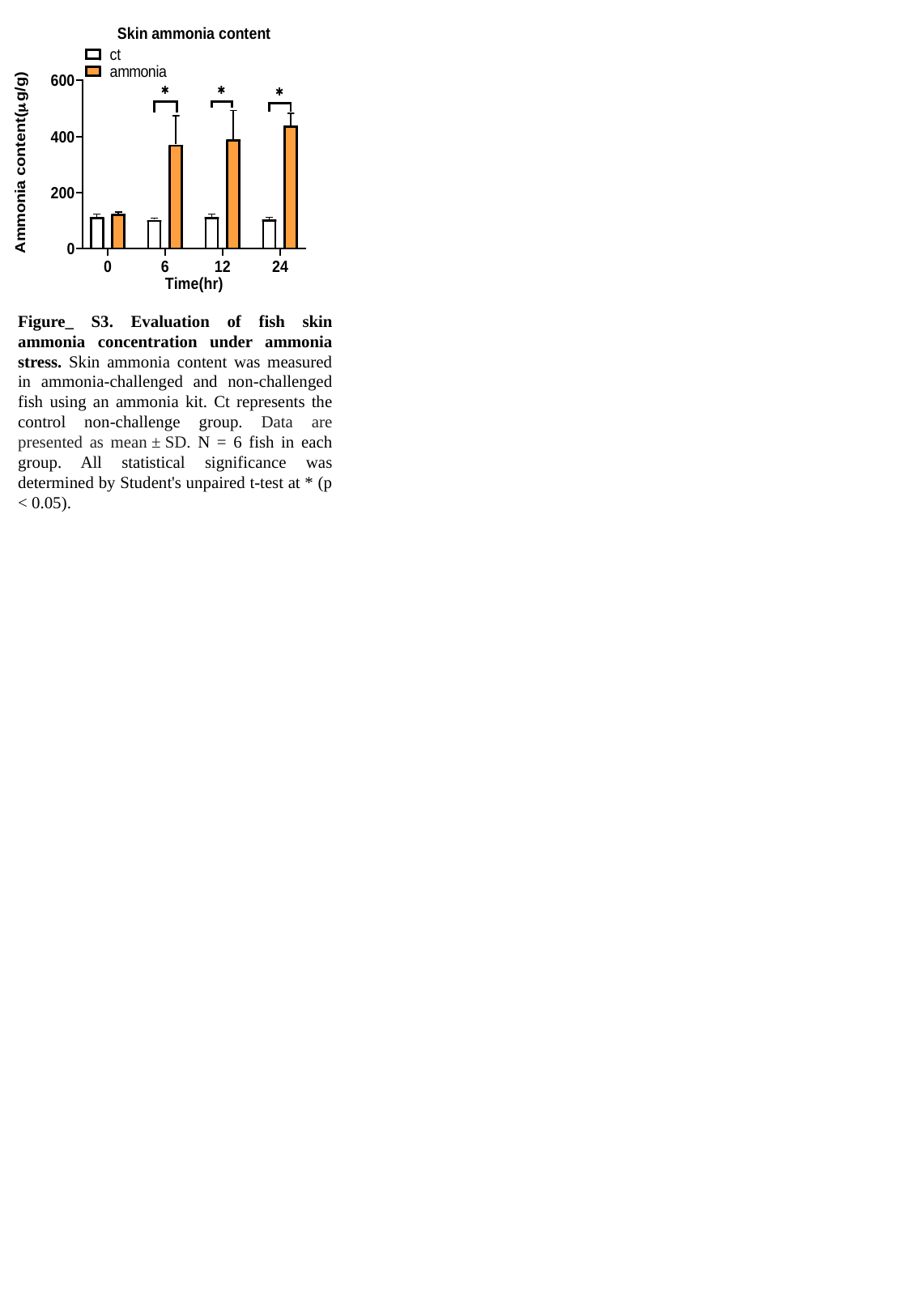

Figure_ S3. Evaluation of fish skin ammonia concentration under ammonia stress. Skin ammonia content was measured in ammonia-challenged and non-challenged fish using an ammonia kit. Ct represents the control non-challenge group. Data are presented as mean ± SD. N = 6 fish in each group. All statistical significance was determined by Student's unpaired t-test at * (p < 0.05).

### Slide 5
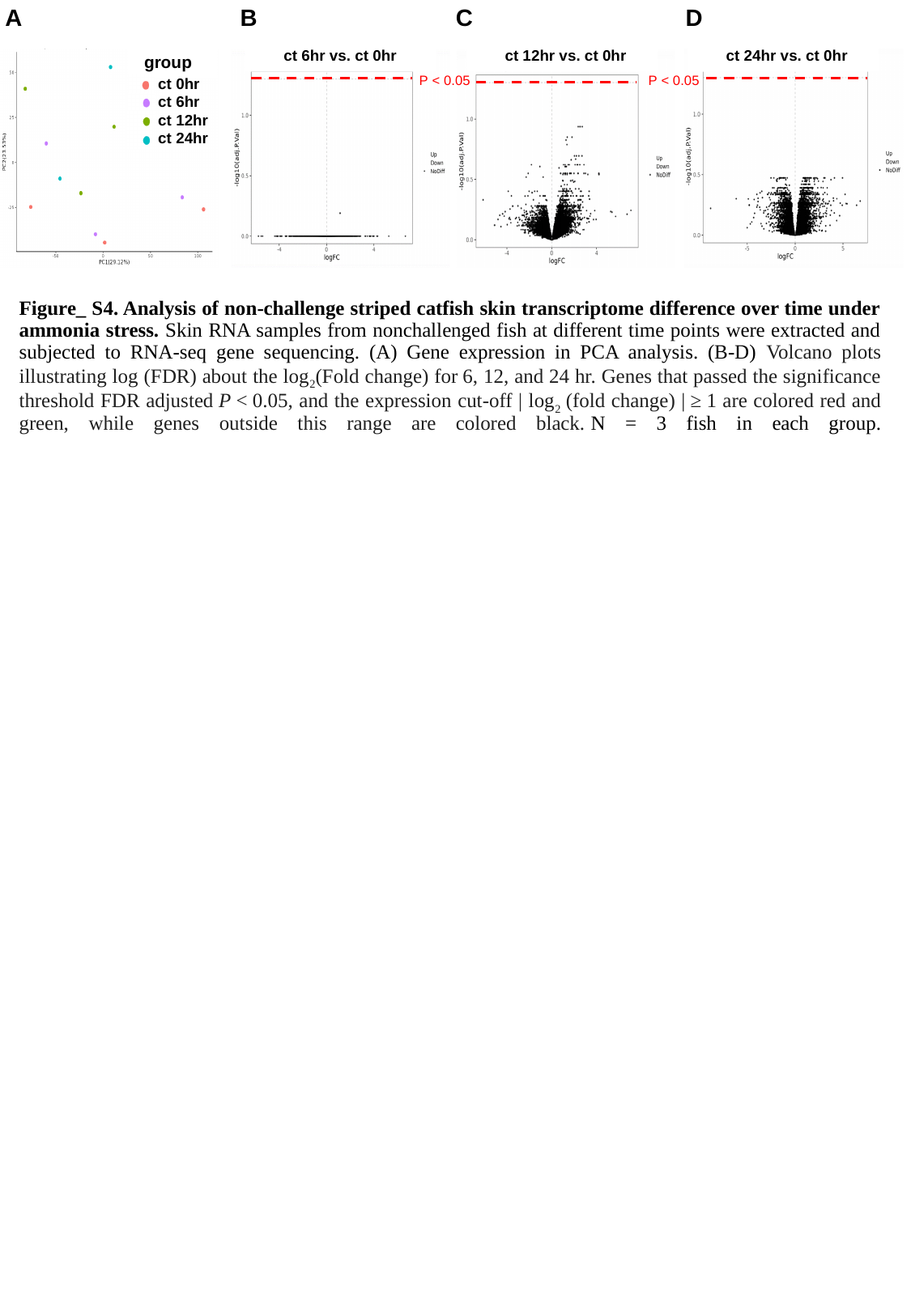

A
B
C
D
 ct 6hr vs. ct 0hr
 ct 12hr vs. ct 0hr
 ct 24hr vs. ct 0hr
 group
P < 0.05
P < 0.05
ct 0hr
ct 6hr
ct 12hr
ct 24hr
#### Figure_ S4. Analysis of non-challenge striped catfish skin transcriptome difference over time under ammonia stress. Skin RNA samples from nonchallenged fish at different time points were extracted and subjected to RNA-seq gene sequencing. (A) Gene expression in PCA analysis. (B-D) Volcano plots illustrating log (FDR) about the log_2_(Fold change) for 6, 12, and 24 hr. Genes that passed the significance threshold FDR adjusted P < 0.05, and the expression cut-off | log_2_ (fold change) | ≥ 1 are colored red and green, while genes outside this range are colored black. N = 3 fish in each group.

### Slide 6
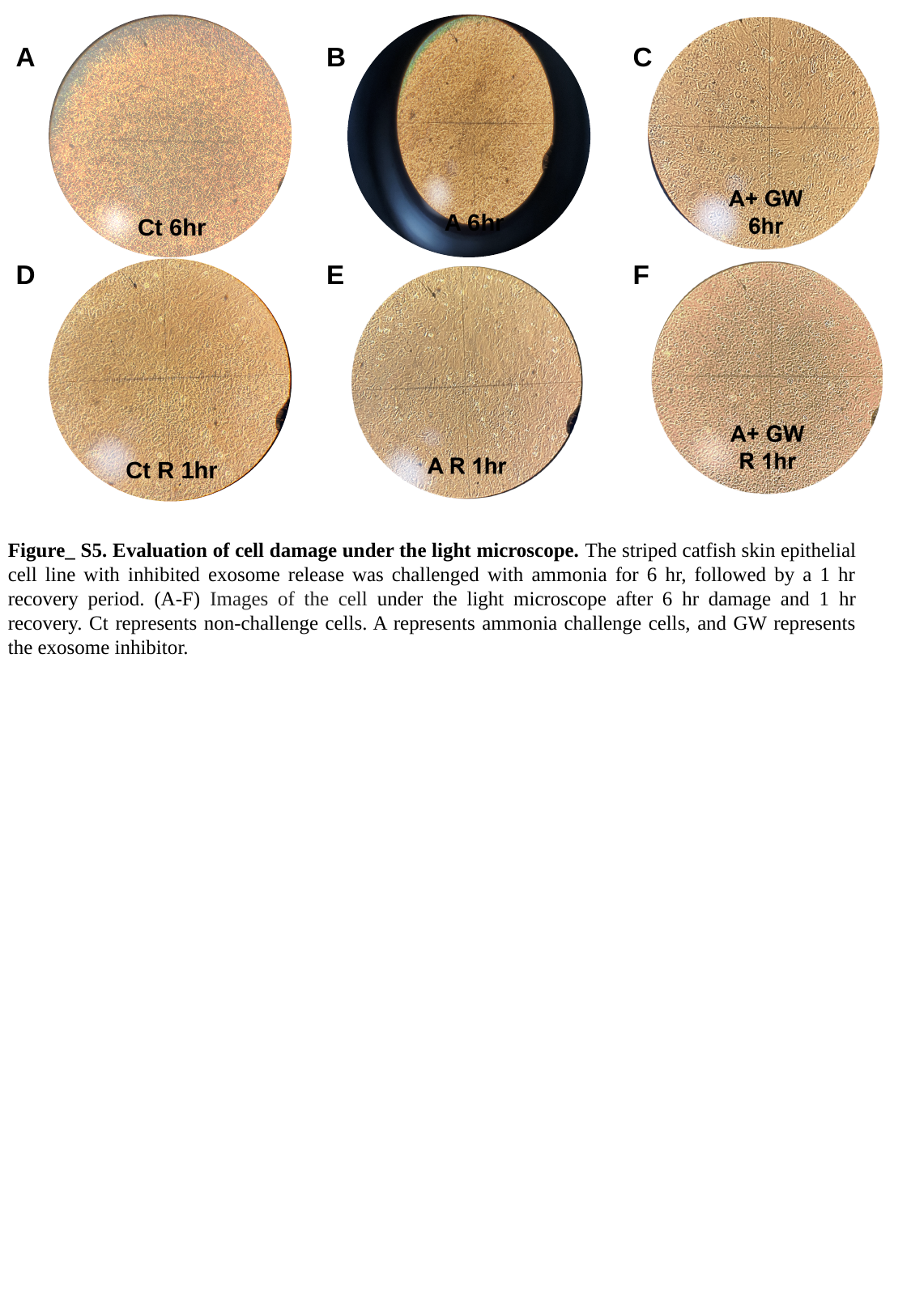

A
B
C
A 6hr
Ct 6hr
D
E
F
Ct R 1hr
Figure_ S5. Evaluation of cell damage under the light microscope. The striped catfish skin epithelial cell line with inhibited exosome release was challenged with ammonia for 6 hr, followed by a 1 hr recovery period. (A-F) Images of the cell under the light microscope after 6 hr damage and 1 hr recovery. Ct represents non-challenge cells. A represents ammonia challenge cells, and GW represents the exosome inhibitor.

### Slide 7
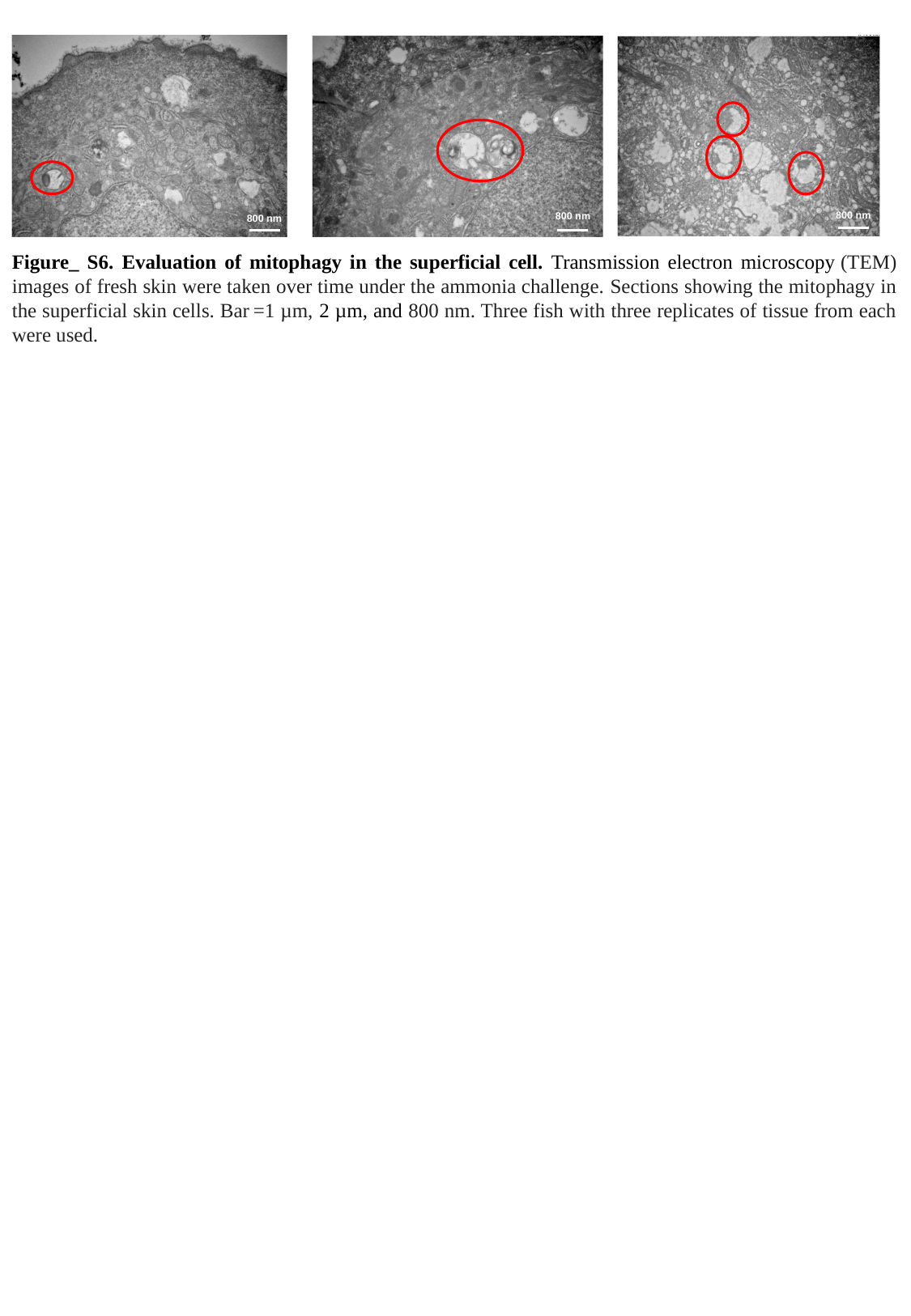

800 nm
800 nm
800 nm
Figure_ S6. Evaluation of mitophagy in the superficial cell. Transmission electron microscopy (TEM) images of fresh skin were taken over time under the ammonia challenge. Sections showing the mitophagy in the superficial skin cells. Bar =1 µm, 2 µm, and 800 nm. Three fish with three replicates of tissue from each were used.

### Slide 8
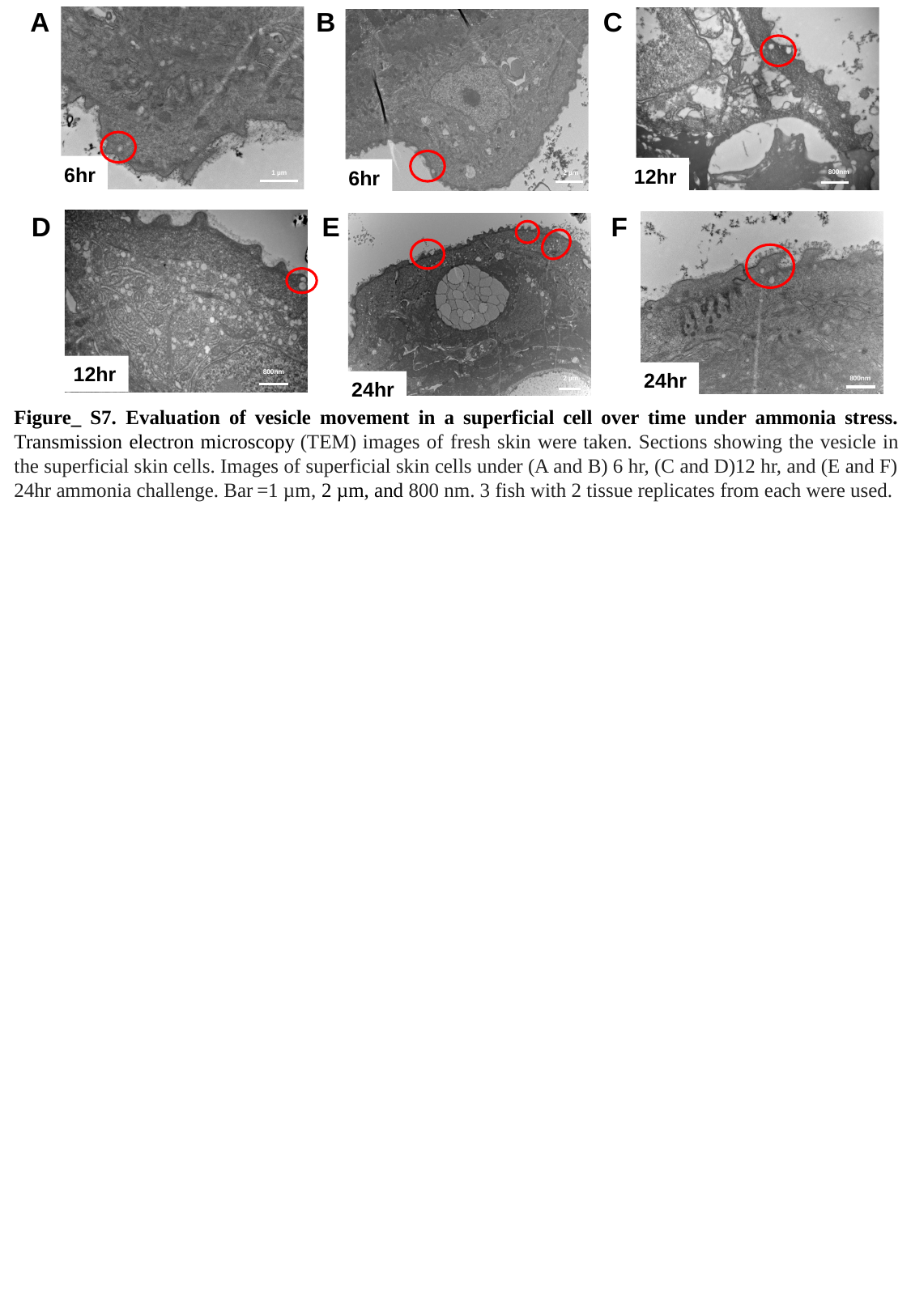

A
B
C
6hr
1 µm
12hr
6hr
2 µm
800nm
D
E
F
12hr
800nm
24hr
800nm
24hr
2 µm
Figure_ S7. Evaluation of vesicle movement in a superficial cell over time under ammonia stress. Transmission electron microscopy (TEM) images of fresh skin were taken. Sections showing the vesicle in the superficial skin cells. Images of superficial skin cells under (A and B) 6 hr, (C and D)12 hr, and (E and F) 24hr ammonia challenge. Bar =1 µm, 2 µm, and 800 nm. 3 fish with 2 tissue replicates from each were used.

### Slide 9
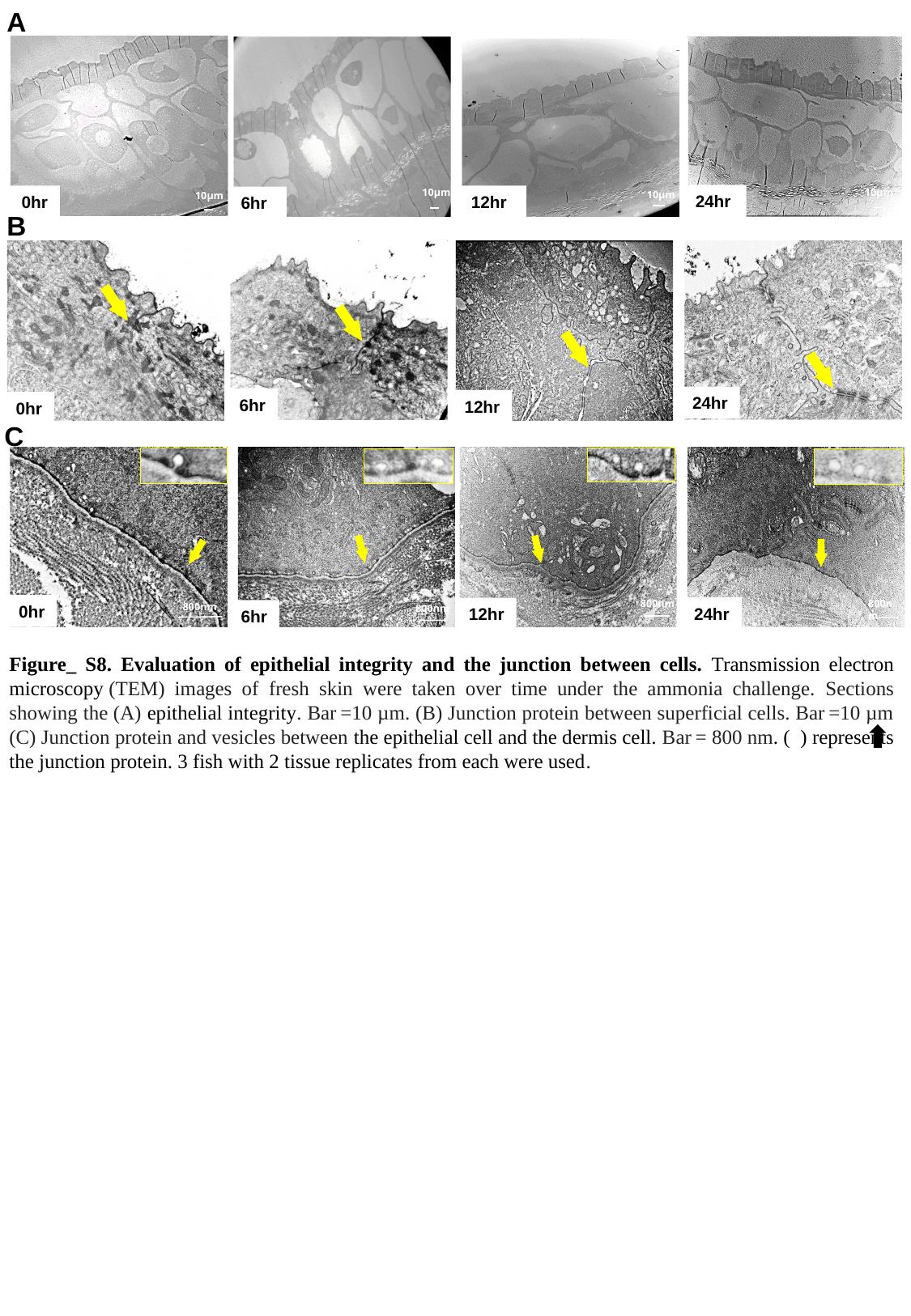

A
10μm
10μm
10μm
24hr
0hr
12hr
6hr
10μm
B
24hr
6hr
12hr
0hr
C
800nm
800nm
0hr
800nm
800nm
24hr
12hr
6hr
Figure_ S8. Evaluation of epithelial integrity and the junction between cells. Transmission electron microscopy (TEM) images of fresh skin were taken over time under the ammonia challenge. Sections showing the (A) epithelial integrity. Bar =10 µm. (B) Junction protein between superficial cells. Bar =10 µm (C) Junction protein and vesicles between the epithelial cell and the dermis cell. Bar = 800 nm. ( ) represents the junction protein. 3 fish with 2 tissue replicates from each were used.
